# Robust decomposition of cell type mixtures in spatial transcriptomics

**DOI:** 10.1101/2020.05.07.082750

**Authors:** Dylan M. Cable, Evan Murray, Luli S. Zou, Aleksandrina Goeva, Evan Z. Macosko, Fei Chen, Rafael A. Irizarry

**Affiliations:** Department of Electrical Engineering and Computer Science, MIT, Cambridge, MA, 02139; Broad Institute of Harvard and MIT, Cambridge, MA, 02142; Department of Data Sciences, Dana-Farber Cancer Institute, Boston, MA, 02215; Department of Biostatistics, Harvard University, Boston, MA, 02115; Department of Psychiatry, Massachusetts General Hospital, Boston, MA, 02114

## Abstract

Spatial transcriptomic technologies measure gene expression at increasing spatial resolution, approaching individual cells. However, a limitation of current technologies is that spatial measurements may contain contributions from multiple cells, hindering the discovery of cell type-specific spatial patterns of localization and expression. Here, we develop Robust Cell Type Decomposition (RCTD, https://github.com/dmcable/RCTD), a computational method that leverages cell type profiles learned from single-cell RNA sequencing data to decompose mixtures, such as those observed in spatial transcriptomic technologies. Our approach accounts for platform effects introduced by systematic technical variability inherent to different sequencing modalities. We demonstrate RCTD provides substantial improvement in cell type assignment in Slide-seq data by accurately reproducing known cell type and subtype localization patterns in the cerebellum and hippocampus. We further show the advantages of RCTD by its ability to detect mixtures and identify cell types on an assessment dataset. Finally, we show how RCTD’s recovery of cell type localization uniquely enables the discovery of genes within a cell type whose expression depends on spatial environment. Spatial mapping of cell types with RCTD has the potential to enable the definition of spatial components of cellular identity, uncovering new principles of cellular organization in biological tissue.

## Introduction

Tissues are composed of diverse cell types and states whose spatial organization governs interaction and function. Recent advances in spatial transcriptomics technologies [1–3] have enabled high throughput collection of RNA-sequencing coupled with spatial information in biological tissues. Using such technologies to spatially map cell types is fundamental to our understanding of tissue structure. In particular, knowledge of spatial localization of specific cellular subtypes remains incomplete and laborious to obtain [4, 5].

Spatial transcriptomics technologies have the potential to elucidate interactions between cellular environment and gene expression, augmenting our knowledge of healthy functions and disease states of tissues. Spatial transcriptomics data is composed of gene expression counts for each of the spatial measurement locations, here referred to as *pixels*, that tile a two dimensional surface. A common task of interest is identifying genes with expression varying across space. Current computational methods search for spatial patterns in gene expression without stratifying by cell type [6–8]. However, much of the variation detected by these methods may be driven by varying cell type composition across the spatial landscape, since single-cell RNA sequencing (scRNA-seq) studies have revealed that cell type can explain a majority of the variation within a population of cells [9, 10]. It is therefore necessary to consider cell type information when searching for spatial gene expression patterns.

Assignment of cell types is analytically challenging, even for high-resolution approaches such as Slide-seq, due to the fact that although pixel resolution can approach the size of mammalian cells (e.g. Slide-seq, 10 microns) [11], fixed pixel locations may overlap with multiple cells. As a result, gene expression measurements at a single pixel may be the result of a mixture of multiple cell types. Currently, the most widely used approach to identifying cell types relies on unsupervised clustering [12]; however, this approach does not allow for the possibility of cell type mixtures. A fundamental challenge is thus to correctly identify these mixture pixels as a combination of multiple cell types, permitting a more complete characterization of the spatial localization of cell types in spatial transcriptomics.

Here, we introduce Robust Cell Type Decomposition (RCTD), a supervised learning approach to decompose RNA sequencing mixtures into single cell types, enabling assignment of cell types to spatial transcriptomic pixels. Specifically, we leverage annotated scRNA-seq data to define cell type-specific profiles for the cell types expected to be present in the spatial transcriptomics data. Supervised cell type assignment methods have achieved high accuracy in scRNA-seq [12,13], but they are not designed for mixtures of multiple cell types. RCTD fits a statistical model that estimates mixtures of cell types at each pixel.

A pertinent challenge for supervised cell type learning is what we term *platform effects*: the effects of technology-dependent library preparation on the capture rate of individual genes between sequencing platforms. We show that if these platform effects are not accounted for, supervised methods are unlikely to succeed since systematic technical variability dominates relevant biological signals [14]. These effects have been previously found in comparisons between single-cell and single-nucleus RNA-seq on the same biological sample [15], where it has been shown that e.g. nucleus-localized genes are enriched in single-nucleus RNA-seq. Here, we demonstrate that platform effects between the scRNA-seq reference and spatial transcriptomics target present a challenge when transferring cell type knowledge to spatial transcriptomics. To enable cross-platform learning in RCTD, we have developed and validated a platform effect normalization procedure.

We demonstrate that RCTD can accurately discover localization of cell types in both simulated and real spatial transcriptomic data. Furthermore, we show that RCTD can detect subtle transcriptomic differences to spatially map cellular subtypes. Finally, we use RCTD to compute expected cell type-specific gene expression, which enables detection of changes in gene expression based on the spatial environment of a cell. Below, we demonstrate how RCTD learns mixtures of cell types in spatial transcriptomics data, facilitating quantification of the effect of spatial position and local cellular environment on gene expression within a cell type.

## Results

### Spatial transcriptomics presents novel challenges: cell type mixtures and platform effects

Spatial transcriptomics pixels source RNA from multiple, rather than single, cells creating a novel challenge for cell type learning. In Slide-seq cerebellum data, we found that the most widely used approach for scRNA-seq cell type identification, unsupervised clustering [12], incorrectly classifies cell types that colocalize spatially but are not similar transcriptomically. For example, Bergmann and Purkinje cells spatially colocalize to the same layer, resulting in a population of pixels that possess marker genes from both cell types (Figure 1a). The most likely explanation for this observation is that these pixels contain two or more cells of different types, but unsupervised clustering assigns these *doublet* pixels to just one cell type. Moreover, this approach predicts granule cells not exclusively in the granular layer, with many cells incorrectly found inside the molecular layer and oligodendrocyte layer (Figure 1b-c, Supplementary Figure 1).

**Figure 1:**
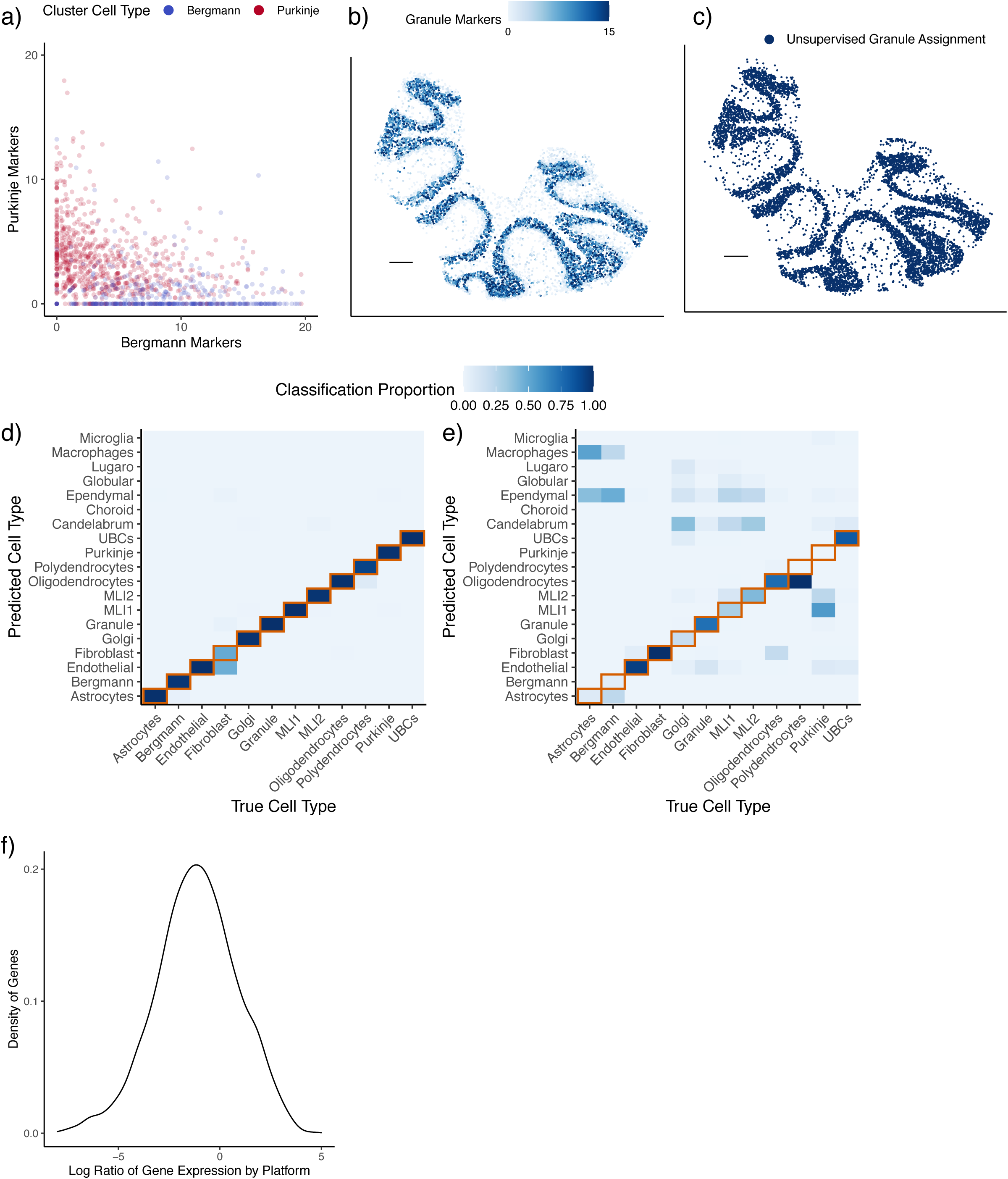
Spatial transcriptomics data presents novel challenges for cell type learning. a) Expression of Bergmann and Purkinje marker genes for pixels colored by unsupervised clustering cell type assignment within a Slide-seq cerebellum dataset. The e.g. Bergmann markers axis is the sum of the expression (counts per 500) of Bergmann differentially expressed genes. b) Expression (counts per 500) of granule marker genes in Slide-seq. Scale bar: 250 microns. c) Spatial plot of granule cells identified by unsupervised clustering. Scale bar: 250 microns. d) Confusion matrix of true vs predicted cell types within training dataset (single-nucleus RNA-seq) by non-negative least squares regression. Color represents the proportion of the cell type on the *x*-axis classified as the cell type on the *y*-axis. The diagonal representing ground truth is boxed in red. e) Confusion matrix of cell type predictions across platforms using non-negative least squares regression trained on single-nucleus RNA-seq, tested on single-cell RNA-seq. Same color scale as (d). f) Density plot, across genes, of measured platform effects between cerebellum single-cell RNA-seq and single-nucleus RNA-seq. The platform effect is defined as the log2 ratio of average gene expression between platforms.

An additional challenge, platform effects, arises in applying supervised learning, in which scRNA-seq cell type profiles are leveraged to classify spatial transcriptomic cell types. For instance, a standard supervised learning approach trained on an assessment single-nucleus RNA-seq cerebellum dataset with known cell types obtained much higher accuracy in the training platform than the testing platform, a single-cell RNA-seq cerebellum dataset (Figure 1d-e). This difference is explained by the presence of *platform effects*: the fact that gene expression changes multiplicatively between single-nucleus and single-cell RNA-seq (Figure 1f). NMFreg, a supervised cell-type mixture assignment algorithm previously developed for Slide-seq, also does not account for platform effects. Testing on the Slide-seq cerebellum dataset, NMFreg assigned a minority (24.8% out of *n* = 11626) of pixels confidently to cell types and mislocalized broad cell type classes (Supplementary Figure 2).

### Robust Cell Type Decomposition enables cross-platform detection of cell type mixtures

To address these challenges, RCTD accounts for platform effects while using a scRNA-seq reference to decompose each spatial transcriptomics pixel into a mixture of individual cell types. RCTD first calculates the mean gene expression profile of each cell type within the annotated scRNA-seq reference (Figure 2a). Next, RCTD fits each spatial transcriptomics pixel as a linear combination of individual cell types, yielding a spatial map of cell types. RCTD takes as input RNA-sequencing counts for each pixel and assumes an unknown mixture of multiple cells (Figure 2a). Each cell type contributes an unobserved proportion of counts to each gene. RCTD estimates the proportion of each cell type for each pixel by fitting a statistical model where, for each pixel *i* and gene *j*, the observed gene counts *Y*_*i,j*_ are assumed to be Poisson-distributed with expected rate determined by *λ*_*i,j*_, a mixture of *K* cell type expression profiles, multiplied by the pixel’s total transcript count, *N*_*i*_:

**Figure 2:**
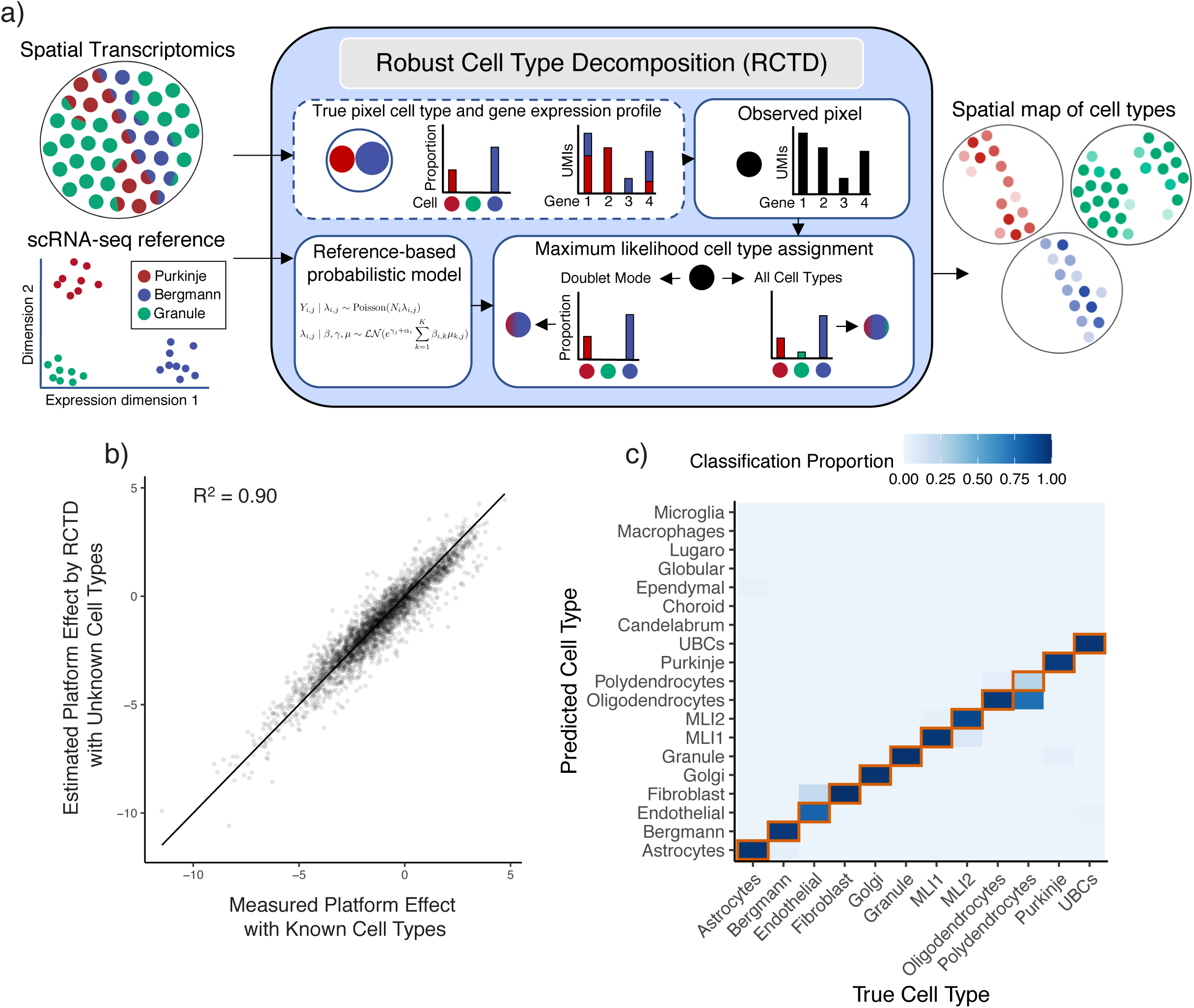
Robust Cell Type Decomposition enables cross-platform learning of cell types. a) Left: RCTD inputs: a scRNA-seq dataset, annotated by cell type, and a spatial transcriptomics dataset with unknown cell types. Middle: RCTD uses a scRNA-seq reference-based probabilistic model to predict cell types on a single pixel containing a mixture of two cell types (e.g. Bergmann/Purkinje), with unknown cell type proportions. RCTD predicts the maximum like-lihood cell type proportions. In *doublet mode*, RCTD constrains each pixel to contain at most two cell types; alternatively, RCTD can estimate the best fit at a pixel using all cell types. Right: RCTD outputs a spatial map of cell types, with opacity representing the inferred cell type proportion. b) Scatter plot of measured vs predicted platform effect (by RCTD) for each gene between the single-cell and single-nucleus cerebellum datasets. Line is the identity line. Measured platform effect is calculated as the log2 ratio of average gene expression between platforms. c) Confusion matrix for RCTD’s performance on cross-platform (trained on single-nucleus RNA-seq, tested on single-cell RNA-seq) cell type assignments for single cells. Color represents the proportion of the cell type on the *x*-axis classified as the cell type on the *y*-axis. The diagonal representing ground truth is boxed in red. All: RCTD was trained on the single-nucleus RNA-seq cerebellum dataset and tested on a dataset of simulated mixtures of single cells from a single-cell RNA-seq cerebellum dataset.

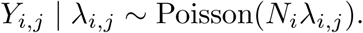

To account for platform effects and other sources of natural variability, such a spatial variability, we assume *λ*_*i,j*_ is a random variable defined by

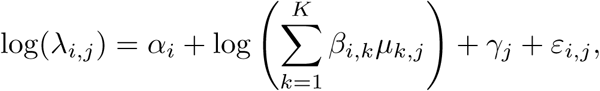

with *µ*_*k,j*_ the mean gene expression profile for cell type *k, α*_*i*_ a fixed pixel-specific effect, *γ*_*j*_ a gene-specific platform random effect and *ε*_*i,j*_ a random effect to account for gene-specific overdispersion.

We use maximum likelihood estimation to infer the cell type proportions, *β*_*i,k*_, indicating which cell types are present in each pixel (see **Methods** for details). RCTD may be used without constraining the number of cell types per pixel or with what we refer to as *doublet mode*, which searches for the best fitting one or two cell types per pixel (see **Methods** for details). In particular, we refer to pixels as *singlets* if they contain only one cell type and *doublets* if they contain two cell types. Doublet mode may mitigate overfitting if mixtures of three or more cell types are expected to be rare, as in Slide-seq [11].

Because gene-specific platform effects are not observable from the raw data, we developed a procedure to estimate platform effects between sequencing platforms with RCTD (**Methods**, Supplementary Table 1). Training RCTD on the single-nucleus RNA-seq cerebellum reference and testing on the single-cell RNA-seq cerebellum dataset, we validated that our approach is able to reliably recover the platform effects (*R*^2^ = 0.90) (Figure 2b). After normalizing cell type profiles for platform effects, RCTD achieved high cross-platform single-cell classification accuracy (89.5% of *n* = 3960 cells) (Figure 2c). Transcriptomically similar cell types, e.g. oligodendrocytes/polydendrocytes, accounted for most of the remaining errors (91.8% of *n* = 415 errors).

To evaluate RCTD’s ability to detect and decompose mixtures in spatial transcriptomics data in the presence of platform effects, we trained RCTD on the single-nucleus RNA-Seq (snRNA-seq) cerebellum reference (Supplementary Figure 3), and tested on a dataset of doublets simulated as computational mixtures of single cells with known cell types in the scRNA-seq dataset (See **Methods** for details). By varying the true underlying cell type proportion, we observed that RCTD correctly classified singlets (92.8% *±* 0.4% s.e.) and doublets (82.8% *±* 0.3% s.e.) with high accuracy (Figure 3a). Additionally, RCTD identified each cell class present on each doublet with 91.9% accuracy on confident calls (**Methods**, *±* 9.4% s.d. across 132 cell type pairs) (Figure 3b). Finally, RCTD accurately estimated the proportion of each cell type on the sample with 12.8% RMSE (*±* 6.9% s.d. across 66 cell type pairs) (Figure 3c-d). These technical validations show that RCTD can accurately learn cell type information in a dataset with mixtures of single cells.

**Figure 3:**
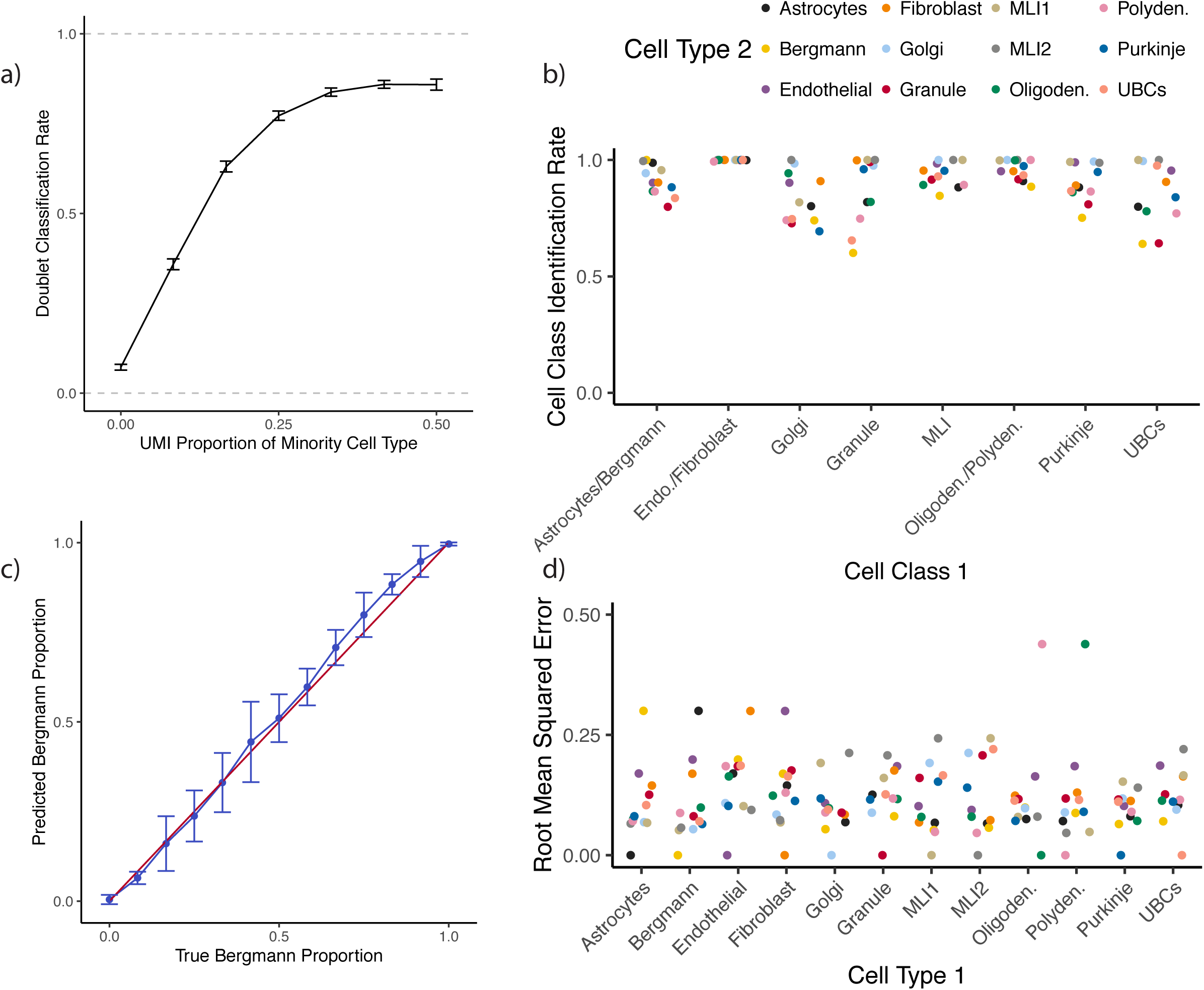
RCTD performs cross-platform detection and decomposition of doublets. a) Rate of doublet classification by RCTD on simulated mixtures of single cells, with 95% confidence intervals. The x-axis represents the true proportion of UMIs sampled from the minority cell type, ranging from 0% (true singlet) to 50% (equal proportion doublet) (1980 ≤ *n* ≤ 3860 simulations per condition). b) On simulated doublets of cell type 1 and cell type 2, the percentage of confident calls by RCTD that correctly identify the cell class of cell type 1, where cell classes group four pairs of transcriptomically similar cell types based on a previous dendrogram [17] (polydendrocytes/oligodendrocytes, MLI1/MLI2, Bergmann/astrocytes, endothelial/fibroblasts). Column represents cell type 1, and color represents cell type 2. c) On simulated Bergmann-Purkinje doublets, predicted Bergmann proportions by RCTD. The *x*-axis represents the true proportion of UMIs sampled from the Bergmann cell. The red line is the identity line, and the blue line is the average and standard deviation (*n* = 30 simulations per condition) of RCTD’s prediction. d) For each pair of cell types, root mean squared error (RMSE) of predicted vs true cell type proportion (as in (c)) by RCTD on simulated doublets (*n* = 390 simulations per cell type pair). Column represents cell type 1, and color represents cell type 2.

### RCTD localizes cell types in spatial transcriptomics data

We next applied RCTD to assign and decompose cell types in spatial transcriptomics data. We first applied RCTD to localize cell types in the mouse cerebellum, using a single-nucleus RNA-seq (snRNA-Seq) reference for training, and a Slide-seqV2 dataset collected on the adult mouse cerebellum as the target. RCTD confidently classified a majority (86.9%, out of *n* = 11626) of pixels, and the resulting cell type calls are consistent with the spatial architecture of the cerebellum (Figure 4a) [16]. To assess performance, we first considered Purkinje/Bergman cells, two cell types which are spatially co-localized in the cerebellum. We found that RCTD’s singlet pixels assigned to Purkinje or Bergmann cell types do not possess markers of the other cell type (Figure 4b). Moreover, pixels predicted as doublets contained marker genes of both Bergmann and Purkinje cells, with estimated cell type proportion correlating with marker gene ratio (Figure 4c). We next observed that RCTD correctly localized molecular layer interneurons to the molecular layer [17], granule cells to the granular layer, and oligodendrocytes to the white matter layer [16], predictions further supported by the spatial correspondence between RCTD’s assignments and the marker genes of each cell type (Figure 4d, Supplementary Figure 4). Next, to validate RCTD’s ability to correctly localize doublets, we leveraged the layered organization of the cerebellum (Figure 4e) [16]. RCTD finds doublets within a layer and between adjacent layers, but rarely between spatially separated layers (Figure 4f).

**Figure 4:**
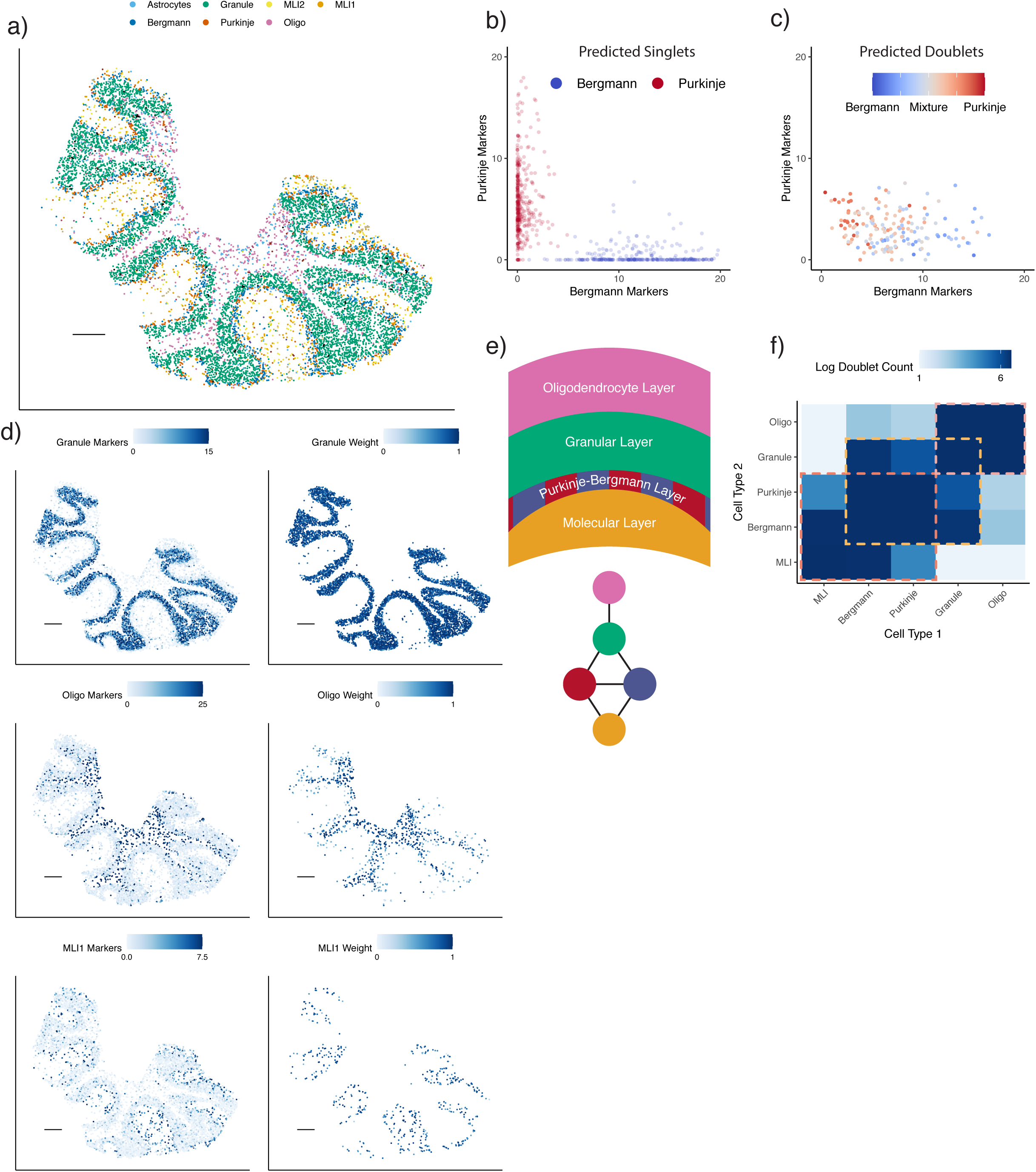
RCTD applied to cell type learning in cerebellum Slide-seq dataset. a) RCTD’s spatial map of cell type assignments in the cerebellum. Out of 19 cell types, the seven most common appear in the legend (individual cell types displayed in Supplementary Figure 4). b) Analogous to (1a), expression of Bergmann and Purkinje marker genes for RCTD’s predicted single cells within a Slide-seq cerebellum dataset (colored by cell type assignment). The e.g. Bergmann markers axis is the sum of the expression (counts per 500) of Bergmann differentially expressed genes. c) Expression of Bergmann and Purkinje marker genes for doublet pixels predicted by RCTD, colored by predicted cell type proportion. d) Predicted spatial localization of cell types by RCTD for granule, oligodendrocytes, and molecular layer interneurons 1 (MLI1). Left: summed expression (counts per 500) (represented by color) of cell type-specific marker genes. Right: predicted spatial locations of each cell type, with color representing predicted cell type proportion. e) (Top) Schematic of spatial cell type organization within the cerebellum [16]. (Bottom) Connectivity graph of cell types that are likely to spatially colocalize. Cell types are colored as in (a). f) Frequency of doublets identified by RCTD between each pair of cell types. Color represents log2 scale counts. Dotted boxes represent communities anatomically expected to exhibit spatial co-localization. Diagonal represents prevalence of singlets. Color bar range: 2 to 100 counts.

### RCTD discovers spatial localization of cellular subtypes

Next, we tested the ability of RCTD to profile the spatial localization of cellular subtypes, recently defined by large-scale transcriptomic analyses [18], for which there is limited knowledge of spatial position in their resident tissues. To this end, we validated RCTD’s ability to classify previously defined [18] subtypes of interneurons in the hippocampus (**Methods**). We first used RCTD to spatially annotate cell types in Slide-seq data of the mouse hippocampus (Figure 5a), training on a scRNA-seq hippocampus dataset [18]. We found that RCTD correctly localizes hippocampal cell types (Supplementary Figure 5, 6). We also validated RCTD’s ability to localize hippocampal cell types in a Visium spatial transcriptomics dataset (Supplementary Figure 6, 7) [2]. We then observed spatial clustering of pixels assigned to the broad class of interneurons (Figure 5b), which we inferred to be derived from large, single interneuron cells [4], an inference supported by histological examination [19]. (Supplementary Figure 8). Consequently, we tested RCTD’s performance in assigning pixels within a cluster to the same interneuron subclass and found high agreement (97.1% *±* 0.09% s.e.) of coarse subclass classification between confident pixels within the same spatial cluster (Figure 5c, **Methods**). Additionally, we found that the spatial localization of the Basket/OLM subclass coincides with expression of *Sst*, a differentially expressed gene for this subclass (Figure 5d). Finally, we used RCTD to assign each spatial cluster to one of 27 transcriptomically defined interneuron subtypes, confidently classifying the majority of interneuron pixels (Figure 5e). Localizations of known subtypes, such as CA1-Lacunosum, which appears in the stratum lacunosum-moleculare (SLM) layer of the CA1 [20], and OLM, which appears primarily in the stratum oriens (SO) [21], agree with known anatomy. We conclude that RCTD enables the identification of spatial locations of cellular subtypes in spatial transcriptomics data.

**Figure 5:**
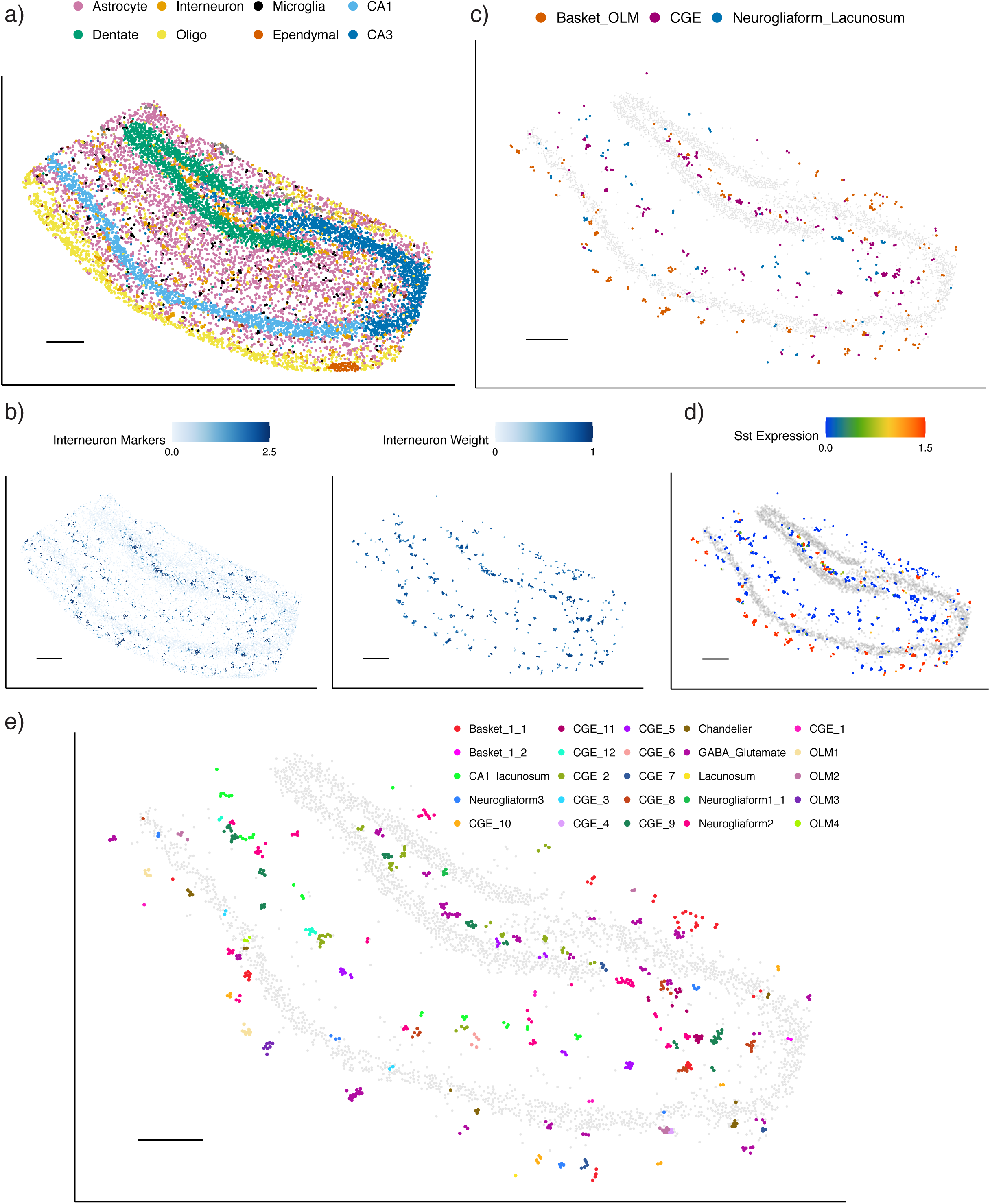
RCTD maps cell types and subtypes in Slide-seq hippocampus. a) RCTD’s spatial map of predicted cell types in the hippocampus. Out of 17 cell types, the 8 most common appear in the legend (individual cell types displayed in Supplementary Figure 5). b) Predicted spatial localization of interneuron cell types by RCTD. Left: normalized expression (represented by color, counts per 500) of marker genes. Right: predicted spatial locations of interneurons, with color representing predicted cell type proportion. c) Predicted confident assignments of interneuron pixels by RCTD to 3 classes of interneuron sub-types, plotted in space. Color indicates predicted subclass. d) Expression (counts per 500) of the *Sst* gene in interneurons identified by RCTD. e) RCTD’s confident assignment of spatial clusters to 27 interneuron subtypes (25/27 subtypes assigned). All scale bars 250 microns. Grey circles represent location of CA1, CA3, and dentate gyrus excitatory neurons for reference.

**Figure 6:**
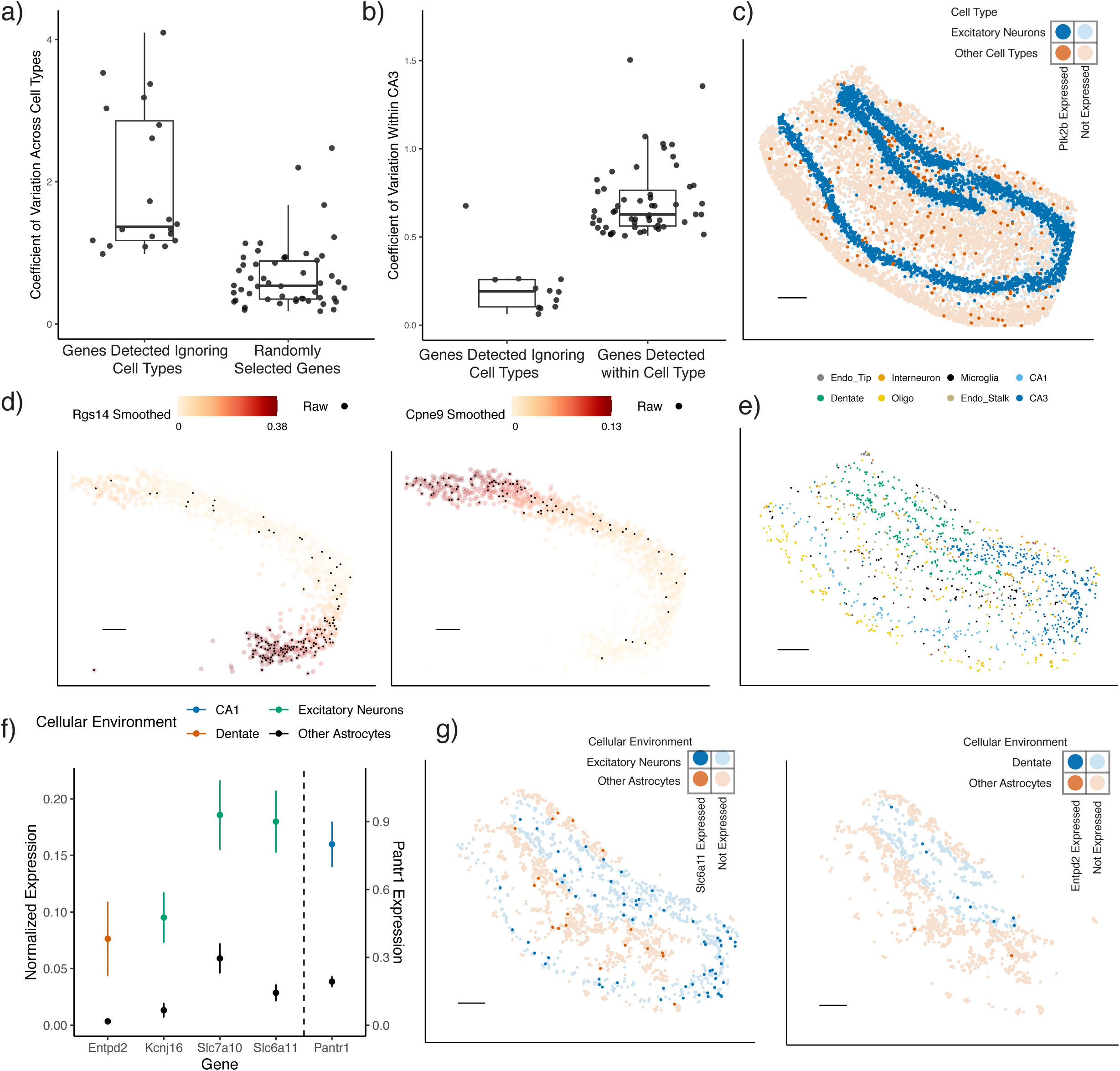
RCTD enables detection of cell type-specific spatial patterns of gene expression. a) Boxplot of coefficient of variation of genes across cell types in the hippocampus single-cell RNA-seq reference. Spatially variable genes were selected for large spatial autocorrelation in the Slide-seq hippocampus, without considering cell type. For reference, 50 randomly selected genes are shown. b-g) Analysis on Slide-seq hippocampus data b) Boxplot of the coefficient of variation in gene expression within CA3 cells identified by RCTD. (Left): Spatially variable genes selected for large spatial autocorrelation in the hippocampus, without considering cell type. (Right): Using RCTD’s expected cell type-specific gene expression, genes determined to be spatially variable by applying local regression within the CA3 cell type (*p* ≤ 0.01, *F*-test). c) Bold pixels represent expression of *Ptk2b*, a gene selected to be spatially variable without considering cell type. Blue represents pixels with excitatory neurons (as detected by RCTD), whereas red represents pixels without excitatory neurons. d) Smoothed spatial expression patterns (counts per 500), recovered by local regression, of two genes detected to have large spatial variation within RCTD’s CA3 cells. Individual pixels expressing the gene are colored in black. e) Spatial localization of astrocyte doublets in the hippocampus, detected by RCTD. Color represents the other cell type on the doublet. f) Mean and standard error of RCTD’s expected gene expression (counts per 500) within groups of astrocytes (129 ≤ *n* ≤ 956 cells per condition) classified by their cellular environment (color). (Scale on the right for *Pantr1*, scale on the left for other genes). g) Spatial visualization of genes with environment-dependent expression within astrocytes. Red represents the astrocytes surrounded by other astrocytes, whereas blue represents astrocytes that are surrounded by excitatory neurons (left) or dentate gyrus cells (right). Bold points represent astrocytes expressing *Slc6a11* (left) or *Entpd2* (right). All scale bars 250 microns.

### RCTD enables detection of spatially variable genes within cell type

Previous computational methods search for spatially variable genes without incorporating cell type information [6–8]. However, because cell types are not evenly distributed in space, and different cell types have different expression profiles, this approach will likely lead to confusing cell type marker genes with spatially variable genes. For example, we found that the 20 genes with the highest spatial autocorrelation in the Slide-seq hippocampus (**Methods**) were primarily expressed in only a few cell types, indicating that their spatial variation is partially driven by cell type composition (Figure 6a). After conditioning on cell type, a majority of these genes exhibited small remaining spatial variation (Figure 6b). For example, *Ptk2b* is differentially expressed in excitatory neurons, but does not exhibit any spatial variation that is unexplained by cell type alone (Figure 6c).

Instead, RCTD enables estimation of spatial gene expression patterns within each cell type. After identifying cell types, we used RCTD to compute the expected cell type-specific gene expression for each cell type within each pixel (see **Methods** for details). Using this cell type-specific expected gene expression, we detected genes with large spatial variation within CA3 pyramidal neurons (Figure 6b, *p* ≤ 0.01, permutation *F*-test, Supplementary Table 2). For these genes, we recovered smooth patterns of gene expression over space with locally weighted regression (Figure 6d, see **Methods** for details). In addition to spatially variable genes, RCTD can be used to detect the effect of cellular environment on gene expression. In the hippocampus, RCTD detected astrocyte doublets with many cell types in distinct spatial regions (Figure 6f); we hypothesized that astrocytic transcriptomes could vary based on their cellular environment. We detected genes whose expression within astrocytes depended on co-localization with another cell type (Figure 6g, **Methods**, Supplementary Table 2). For instance, we found that *Entpd2* was enriched in astrocytes colocalizing with dentate neurons (*p* = .025, *z*-test). This is consistent with a prior study that detected a population of astrocyte-like progenitor cells in the dentate expressing *Entpd2* [22]. Moreover, *Slc6a11*, which enables uptake of the GABA neurotransmitter and likely modulates inhibitory synapses [23], was differentially expressed in astrocytes around excitatory neurons (*p <* 10^−6^, *z*-test) [24]. Thus, RCTD enables measurement of the effect of the cellular environment and space on gene expression.

## Discussion

Accurate spatial mapping of cell types and detection cell type-specific spatial patterns of gene expression is critical for understanding tissue organization and function. Here, we introduce RCTD, a computational method for accurate decomposition of spatial transcriptomic pixels into mixtures of cell types, using a single-cell RNA-seq reference normalized for platform effects. RCTD takes as input RNA sequencing counts at each pixel containing an unknown mixture of multiple cells, and predicts the proportion of each cell type on each pixel. RCTD accurately maps cell types, as demonstrated on both a dataset of simulated doublets as well as cerebellum and hippocampus spatial transcriptomics datasets. We additionally demonstrated RCTD’s ability to correctly localize subtypes in a Visium hippocampus spatial transcriptomics dataset, showing that RCTD can be applied broadly to different platforms. We further showed RCTD can spatially localize transcriptomically-defined cellular sub-types of interneurons of the hippocampus. Lastly, we demonstrated that RCTD enables discovery of spatially varying gene expression within cell types in the hippocampus.

As the cost of sequencing diminishes, scRNA-seq datasets are becoming more prevalent and easier to generate [25]. Individual scRNA-seq methods can be more or less similar to a spatial transcriptomics dataset in their platform effects, which can be measured by RCTD. For example, relative to Slide-seq, we found a lower magnitude of platform effects for the single-cell hippocampus reference than for the single-nucleus cerebellum reference. However, since the single-cell sequencing platform of best-available quality can vary for a given tissue, we have designed RCTD to be flexible to choice of reference. We thus anticipate it to be compatible with future scRNA-seq modalities. Furthermore, our method is flexible to the choice of target platform. For example, our procedure for estimating platform effects depends only on merging all pixels into one *pseudo-bulk* measurement. Our method can consequently be applied to estimate platform effects from a scRNA-seq reference to any other sequencing technology, including bulk RNA sequencing, providing a generally-applicable normalization procedure for RNA sequencing. Although motivated by spatial transcriptomics, we expect that RCTD can learn cell types on other non-spatial datasets with single cells or mixtures of multiple cell types [26].

When fine spatial resolution causes localization of three or more cell types to one pixel to be uncommon (e.g. Slide-seq [11]), we recommend using doublet mode of RCTD, which constrains at most two cell types per pixel. Otherwise, RCTD can be used to decompose any number of cell types per pixel (e.g. Visium). Similar in principle to AIC model selection methods [27], doublet mode reduces overfitting by penalizing the number of cell types used, improving RCTD’s statistical power. This concept can be readily extended to triplets and beyond in future work.

A major goal of spatial transcriptomics is understanding the contributions of cell type and cellular environment on cell state. RCTD facilitates the discovery of these effects by computing expected cell type-specific gene expression for each spatial transcriptomics pixel. For instance, we analyzed gene expression within astrocytes to detect astrocytic genes influenced by local cellular environment. There are many drivers of a gene’s dependence on cellular environment: cell-to-cell interactions, regional signalling factors, or cellular history during development. The ability of RCTD to localize cell types uniquely enables high-throughput generation of biologically-relevant hypotheses concerning the effects of space and environment on gene expression. As more spatial transcriptomics datasets are generated, we expect that RCTD will facilitate the discovery of new principles of cellular organization in biological tissue.

## Methods

### Statistical model

Here, we describe the statistical model used to perform Robust Cell Type Decomposition (RCTD) to identify mixtures of cell types. For each pixel *i* = 1, …, *I* in the spatial transcriptomics dataset, we denote the observed gene expression counts as *Y*_*i,j*_ for each gene *j* = 1, …, *J*. We model these counts with the following hierarchical model,

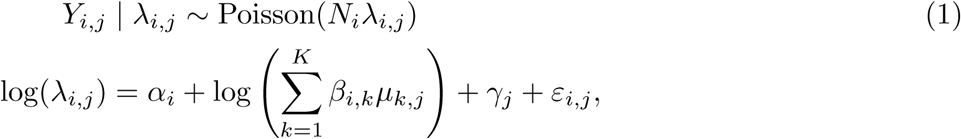

with *N*_*i*_ the total transcript count or number of unique molecular identifies (UMIs) for pixel *i, K* the number of cell types present in our dataset, *α*_*i*_ a fixed pixel-specific effect, *µ*_*k,j*_ the mean gene expression profile for cell type *k* and gene *j, β*_*i,k*_ the proportion of the contribution of cell type *k* to pixel *i, γ*_*j*_ a gene-specific platform random effect and *ε*_*i,j*_ a random effect to account for other sources of variation, such as spatial effects. Data exploration (Figure 1f) supported a Poisson-lognormal mixture, used previously for count data [28]. Thus, we assume *γ*_*j*_ and *ε*_*i,j*_ both follow normal distributions with mean 0 and standard deviation *σ*_*γ*_ and *σ*_*ε*_, respectively. We note that in practice we additionally modify the random effects distributions to include a heavier tail that is robust to outliers (using an approximation to a Cauchy-Gaussian mixture distribution [29]; see supplementary methods for details). The main goal of our analysis is to estimate the *β*_*i,k*_’s, which represent the cell type or cell types present in each pixel *i*, constrained so that 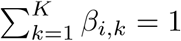 and each *β*_*i,k*_ ≥ 0.

### Fitting the model

Model (1) is a complex model with thousands of parameters (many, *K × J*, of these parameters are introduced by the cell type-specific gene expression profiles). We overcome this challenge by fitting our model using a stepwise approach that includes a supervised learning step for estimating these expression profiles, *µ*_*k,j*_. The steps of our estimation approach are as follows:

1. Supervised estimation of cell type profiles: We use a *reference* dataset, refered to as the *training* dataset, to obtain estimates for the mean gene expression profiles *µ*_*k,j*_. We refer to these estimates as 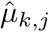, which are then considered fixed in the next steps.
2. Gene filtering: We use the estimated cell type profiles 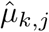 to filter out genes that are unlikely to be informative. We do this by selecting genes that show differential expression across cell types.
3. Platform Effect Normalization: The random effects *γ*_*j*_ account for the unwanted technical variation resulting from gene expression profiles varying across different sequencing platforms. The next step is therefore to estimate *σ*_*γ*_ and predict *γ*_*j*_ for each gene *j*. We denote the prediction of the random effects as 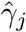, which are then considered fixed in the next step.
4. Robust Cell Type Decomposition: We use the plugin estimates 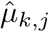 and 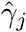 and assume they are fixed. Conditional on these estimates, for each sample *i* and treating *ε*_*i,j*_ as a random effects, we can compute the maximum likelihood estimate (MLE) for *β*_*i,k*_, *α*_*i*_, and *σ*_*ε*_.

Next we describe each of these steps in detail.

### Supervised estimation of cell type profiles

First, we obtain a single-cell RNA-seq reference, which has been previously annotated with cell types. We estimate 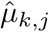 as the average normalized expression of gene *j* within all cells of cell type *k*.

### Gene filtering

Using the estimated cell type expression profiles 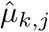, we select differentially expressed genes that will be informative when estimating cell type proportions. For each cell type in the scRNA-seq reference, we select genes with minimum average expression above .0625 counts per 500 and at least 0.5 log-fold-change compared to the average expression across all cell types. Typically, this results in about 5, 000 genes for the platform effect normalization step. These parameters are further increased for the Robust Cell Type Decomposition step, to reduce the set to about 3, 000 genes for computational efficiency.

### Platform effect normalization

Estimating the *β*_*i,k*_ in the presence of the unobserved platform effects *γ*_*j*_ is challenging. However, *γ*_*j*_ can be reliably predicted independently from the other parameters by summarizing the spatial transcriptomics data as a single *pseudo-bulk* measurement 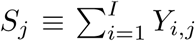. Notice that, conditioned *λ*_*i,j*_, *S*_*j*_ is Poisson distributed with the average 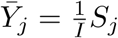 having expectation:

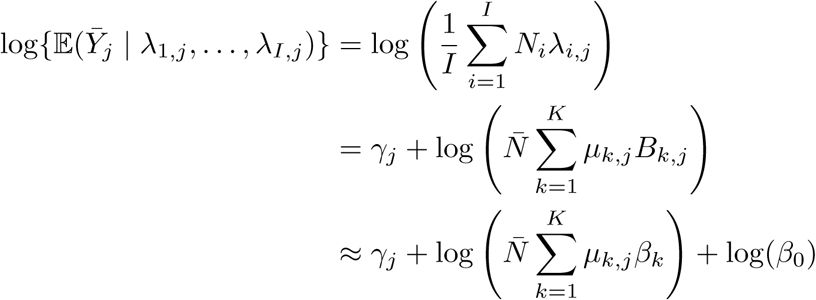

With

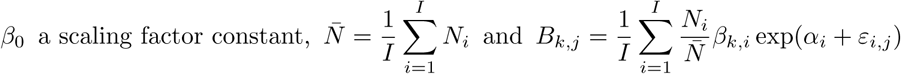

a random variable that is approximately proportional to 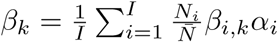, the proportion of cell type *k* in our target dataset:

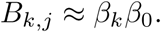

This follows from the fact that 𝔼 (*B*_*k,j*_) = *β*_*k*_*β*_0_, and Var(*B*_*k,j*_) converges to 0 when *I* is large (see supplementary methods for details). By plugging in the 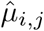 obtained in the first step and treating them as known, we can then obtain the maximum likelihood estimator (MLE) for *β*_0_, the *β*_*k*_’s, and *σ*_*γ*_ and subsequently estimate the platform effects *γ*_*j*_ as 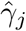.

### Robust Cell Type Decomposition

With 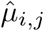 and 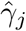 in place, we plug them into equation (1) which we can rewrite as,

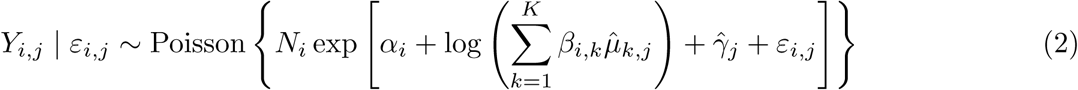

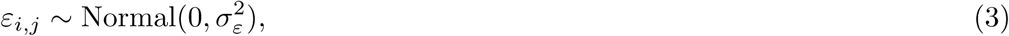

and we obtain the MLE *α*_*i*_, *β*_*i,k*_ and *σ*_*ε*_. The algorithm implemented to find the MLE is in the supplementary methods.

### Cell type identification by model selection

Notice that in the procedure described above, 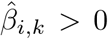 for as many as *K* cell types, implying that pixel *i* is a mixture of several cell types. However, for many spatial transcriptomics technologies, we do not expect more than two cell types per pixel. We therefore implemented a version of our model and estimation procedure that constrains the number of *k*^*t*^s for which *β*_*i,k*_ *>* 0 to two. We refer to this version of method as *doublet* mode. In doublet mode, cell type identification is accomplished using a model selection framework, where we compare likelihoods and penalize the inclusion of an additional features. In this version of our method, we refer to the two possible outcomes as *singlet* and *doublet*.

Specifically, for each cell type *k*, we compute (*k*) as the log-likelihood of the model fit with only cell type *k*, and (*k, ℓ*) as the log-likelihood of the model fit with only cell types *k* and *ℓ*. For each pixel *i* we then define

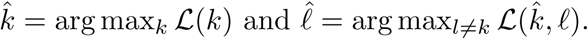

Because we expect many pixels to represent only one cell type, we then used a penalized approach similar to AIC [27] to decide between the two models, using only one cell 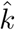 or two 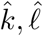. Specifically, we select the model *ℳ* maximizing,

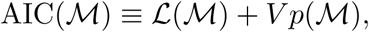

with *p* the number of parameters (cell types) and *V* a penalty weight. In the results presented here, we selected *V* = 25 based on simulation studies.

We then use an ad-hoc approach to classify our selections into either *confident* or *unconfident* in the following way:

1. Consider pairs of cell types (*k, ℓ*) such that 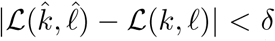. If there exists one such pair such that 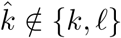 and another (possibly identical) pair where 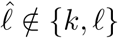, then we assume that we do not have enough information to predict cell types and call this pixel *unconfident*. If this condition does not hold, then we will be *confident* of at least one cell type, 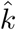 and/or 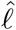, that appears in all such pairs.
2. If condition 1 does not hold, and we select the singlet model, then we call this a *confident singlet*.
3. If condition 1 does not hold, and we select the doublet model, then if there exists a cell type pair {*k, ℓ}* distinct from 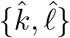 for which 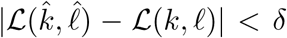, we call this a *unconfident doublet*, otherwise we call this a *confident doublet*.

For the work in this paper, we set *δ* = 10 based on simulation studies.

### Classification of cellular subtypes

We apply the RCTD procedure described above to detect major cell types. But, as mentioned in the results section, recently characterized cellular subtypes have been identified and defined by large-scale transcriptomic analyses [18]. After selecting pixels in which RCTD was confident of the presence of the cell type of interest, we re-ran RCTD on these pixels using a larger set of cellular subtype profiles defined by the reference. During the subtype step of RCTD, we constrained the major cell types appearing on each pixel so be the same as originally detected by RCTD.

For interneurons, we used 27 previously defined [18] interneuron subtypes and hierarchically clustered the log average expression vectors of these subtypes into 3 major subclasses (Supplementary Figure 9). In order to define spatial clusters of Slide-seq interneurons, we hierarchically clustered the points in space and manually split doublets. To classify a set of pixels presumed to comprise the same cell, we selected the subtype maximizing the joint density of these pixels by summing the log-likelihoods.

### Expected cell type-specific gene expression

Once *β* has been estimated by RCTD, we can compute the expected cell type-specific gene expression at each pixel. Specifically, we compute the conditional expectation of *Y*_*i,k,j*_, the expression of gene *j* on pixel *i* from cell type *k* (see supplementary methods for derivation):

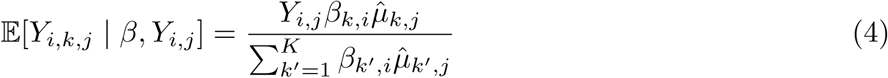

Intuitively, the expected expression of a cell type is proportional to the proportion of the cell type on the pixel and the probability of observing the gene in each cell type. We note that we are only computing the conditional expectation 𝔼 [*Y*_*i,k,j*_ | *β, Y*_*i,j*_], but *Y*_*i,k,j*_ | *β, Y*_*i,j*_ may have large variance for a single pixel, due to sampling noise. Furthermore, this estimate is based on a strong assumption of the model that random effects of gene expression *ε*_*i,j*_ are shared across cell types.

### Collection and processing of scRNA-seq and spatial transcriptomics data

We used publicly available single-cell RNA-seq datasets, which have previously been annotated by cell type. For running RCTD on cerebellum, we trained on a single-nucleus RNA-seq dataset [17]. For training RCTD on hippocampus, and testing (cross-platform) RCTD in cerebellum, we used the DropViz single-cell RNA-seq dataset [18]. The DropViz hippocampus dataset also contained annotations for interneuron subtypes. For marker gene plots, we define a *metagene* for each cell type as the sum of genes that are over-expressed with log-fold-change above 3.

Slide-seq mouse cerebellum data was collected using the Slide-seqV2 protocol, developed and described recently (see supplementary methods for details) [1]. Slide-seqV2 hippocampus and Visium hippocampus data were used from previous studies [1, 2]. Data pre-processing occurred using the Slide-seq tools pipeline [1]. The region of interest was cropped prior to running RCTD, and spatial transcriptomic spots were filtered to have a minimum of 100 UMIs.

### Validation with simulated doublets dataset

We trained RCTD on the cerebellum single-nucleus RNA-seq reference, and tested the model on a dataset of doublets simulated from the single-cell RNA-seq cerebellum dataset. We restricted to 12 cell types that appeared both in the single-nucleus and single-cell reference. In order to simulate a doublet, we randomly chose a cell from each cell type, and sampled a predefined number of UMIs from each cell (total 1, 000). We defined a doublet as containing 25-75% of UMIs for each of the two cell types, whereas a singlet contained 0% or 100%. We defined *doublet classification rate* (Figure 3a) as the ratio of number of predicted doublets to total predicted singlets or doublets. Cell type proportion estimation (Figure 3b, 3c) was measured with RCTD fit using the two cell types present on the simulated doublet. We defined coarser *classes* of cell types (used for Figure 3d) based on a previously defined dendrogram [17]. This resulted in pairing of MLI1/MLI2, Astrocytes/Bergmann, Oligoden./Polyden., and Endothelial/Fibroblast. Cell class identification rate (Figure 3d, top) was calculated on the subset of confidently called cell types.

### Detection of cell type-specific gene expression patterns

After computing expected cell type-specific gene expression, we detected spatially variable genes within a cell type. Genes were filtered for minimum average expression within the scRNA-seq reference of the cell type of interest (.0125 counts per 500, and at least 50% as large as average expression of other cell types). We applied 2D local regression to these genes, and calculated coefficient of variation (CV) of the estimated smooth function. We selected genes with CV ≥ 0.5 and tested the local regression variation with a permutation *F*-test (*p* ≤ 0.01, 99 permutations of spatial locations).

Next, we searched for genes that changed their expression within astrocytes based on co-localization with another cell type. We classified astrocytes as co-localizing with another particular cell type if at least 25% of their neighbors within a 40 micron radius were that cell type. If at least 80% of these neighbors were other astrocytes, the cell was classified as co-localizing with other astrocytes. We filtered for genes in the scRNA-seq reference with average expression log-fold-change of ≥ 2.3 within astrocytes vs. each other cell type. We looked for genes that were differentially expressed depending on the co-localized cell type, testing with a *z*-test (*p <* 0.05).

### Implementation details

RCTD is publicly available as an R package (https://github.com/dmcable/RCTD). The quadratic program that arises in the RCTD optimization algorithm is solved using the quadprog package in R [30]. We used and modified code from the DWLS package to implement sequential quadratic programming for RCTD [31, 32]. Non-negative least squares regression was also implemented as a quadratic program. Unsupervised clustering was performed using the Seurat package, following Seurat’s spatial transcriptomics vignette [33]. Clusters were assigned by their expression of marker genes and spatial localization. Additionally, detection of globally spatially variable genes was accomplished using Seurat’s implementation of Moran’s I. Local regression was accomplished with the loess function. The NMFreg python notebook was used with default parameters (factors = 30) for testing NMFreg.

## Author Contributions

D.M.C., F.C., R.A.I, and E.Z.M. conceived the study; F.C., E.M., and E.Z.M. designed the Slide-seq experiment; E.M. generated the Slide-seq data; D.M.C., R.A.I., and F.C. developed the statistical methods; D.M.C., F.C., R.A.I, and E.Z.M. designed the analysis; D.M.C., R.A.I., F.C, A.G., and L.S.Z. analyzed the data; D.M.C., F.C., R.A.I., E.Z.M., and L.S.Z. wrote the manuscript; all authors read and approved the final manuscript.

### Acknowledgements

We thank Robert Stickels for providing valuable input on the analysis. We thank members of the Chen lab, Irizarry lab, and Macosko lab for helpful discussions. D.C. was supported by a Fannie and John Hertz Foundation Fellowship and an NSF Graduate Research Fellowship. This work was supported by an NIH Early Independence Award (DP5, 1DP5OD024583 to F.C.), the NHGRI (R01, R01HG010647 to E.Z.M. and F.C.), as well as the Schmidt Fellows Program at the Broad Institute and the Stanley Center for Psychiatric Research. R.A.I. was supported by NIH grants R35GM131802 and R01HG005220.

## Code Availability Statement

RCTD is implemented in the open-source R package RCTD, with source code freely available at https://github.com/dmcable/RCTD. Additional code used for analysis in this paper is available at https://github.com/dmcable/RCTD/tree/dev.

## Data Availability Statement

Raw sequence data from this study will be deposited in GEO. Accession codes will be available before publication.

## Conflict of Interest Statement

The authors declare no conflict of interest.

## Notes

https://github.com/dmcable/RCTD

### Competing Interest Statement

The authors have declared no competing interest.

