## Supplementary Methods and Figures for "Robust decomposition of cell type mixtures in spatial transcriptomics"

##### Estimation of cell type means

We estimate $\hat{\mu}_{k,j}$ as the empirical mean of normalized expression of gene $i$ in cell type $j \in J$ within the scRNA-seq reference:

$$\hat{\mu}_{k,j} \equiv \frac{1}{I_k} \sum_{i=1}^{I_k} \frac{Y_{i,k,j}}{N_{i,k}} \tag{1}$$

Here, $I_k$ is the number of cells in reference of cell type $k$, $N_{i,k}$ is the number of UMIs of cell $i$ of cell type $k$, and $Y_{i,k,j}$ is observed counts of gene $j$ in this cell. If we assume there are many cells, then this will be an accurate and unbiased estimator of the true mean $\mu_{k,j}$.

##### Probabilistic model

In this section, we revisit our probabilistic model of spatial transcriptomics, deriving the Poisson sampling model from a more detailed independently-sampled unique molecular identifier (UMI) model. For each UMI, the source cell type is first probabilistically determined based on the cell type mixture proportions. Second, the gene is determined according to the probability vector of that cell type. Formally, for each pixel $1 \leq i \leq I$ and for each read $1 \leq n \leq N_i$, we select cell type $1 \leq \theta_{n,i} \leq K$ and gene $1 \leq z_{n,i} \leq J$ as:

$$P(\theta_{n,i} = k \mid \beta) = \beta_{k,i}, \quad P(z_{n,i} = j \mid \theta_{n,i}, \gamma, \varepsilon) = \delta_{i,\theta_{n,i},j} \tag{2}$$

Here, we define:

$$\log(\delta_{i,k,j}) = \alpha_i + \log(\hat{\mu}_{k,j}) + \gamma_j + \varepsilon_{i,j}, \tag{3}$$

$$\text{where } \gamma_j \sim \text{Normal}(0, \sigma_\gamma^2), \quad \varepsilon_{i,j} \sim \text{Normal}(0, \sigma_\varepsilon^2), \quad \alpha_i \text{ fixed} \tag{4}$$

Next, we define $\lambda_{i,j}$ (predicted gene probability) as:

$$\lambda_{i,j} \equiv P(z_{n,i} = j \mid \gamma, \varepsilon) = \sum_{k=1}^{K} \beta_{i,k} \mu_{k,j} \tag{5}$$

We notice that as a consequence of our model:

$$Y_{i,j} \mid \lambda_{i,j} = \sum_{n=1}^{N_i} \mathbb{I}[z_{n,i} = j] \sim \text{Binom}(N_i, \lambda_{i,j}) \approx \text{Poisson}(N_i \lambda_{i,j}) \tag{6}$$

Thus, we recover the Poisson model fit by RCTD.

##### Platform Effect Normalization

The platform effects $\gamma_i$ can be reliably estimated independently from the other parameters by summarizing the spatial transcriptomics data as a single *pseudo-bulk* measurement $S_j$:

$$S_j \equiv \sum_{i=1}^{I} Y_{i,j} \sim \text{Poisson}(\sum_{i=1}^{I} N_i \lambda_{i,j}) \tag{7}$$

where $S_j$ are observed. Next, we calculate the rate parameter of this Poisson distribution:

$$\begin{aligned} \mathbb{E}[S_j|\gamma] = \sum_{i=1}^{I} N_i \lambda_{i,j} &= \sum_{i=1}^{I} \sum_{k=1}^{K} N_i \beta_{k,i} \delta_{i,k,j} = \sum_{k=1}^{K} \sum_{i=1}^{I} N_i \beta_{k,i} \mu_{k,j} e^{\gamma_j + \alpha_i + \varepsilon_{i,j}} \\ &= e^{\gamma_j} \sum_{k=1}^{K} \mu_{k,j} \sum_{i=1}^{I} N_i \beta_{k,i} e^{\alpha_i} e^{\varepsilon_{i,j}} \\ &= I e^{\gamma_j} \bar{N} \sum_{k=1}^{K} \mu_{k,j} B_{k,j} \\ &\approx I e^{\gamma_j} \bar{N} \sum_{k=1}^{K} \mu_{k,j} \beta_k \beta_0 \end{aligned} \tag{8}$$

Where we have defined $\beta_0 \equiv \mathbb{E}[\varepsilon_{i,j}] = e^{\sigma_\varepsilon^2/2}$ as a fixed scaling constant (using lognormal expectation), $\beta_k = \frac{1}{I} \sum_{i=1}^{I} \frac{N_i}{\bar{N}} \beta_{i,k} \alpha_i$ is the average proportion of cell type $k$, and:

$$\bar{N} = \frac{1}{I} \sum_{i=1}^{I} N_i \text{ and } B_{k,j} = \frac{1}{I} \sum_{i=1}^{I} \frac{N_i}{\bar{N}} \beta_{k,i} \exp(\alpha_i + \varepsilon_{i,j}). \tag{9}$$

$B_{k,j}$ is a random variable that is approximately proportional to $\beta_k$:

$$B_{k,j} \approx \beta_k \beta_0.$$

This follows from the fact that $\mathbb{E}(B_{k,j}) = \beta_k \beta_0$, and $\text{Var}(B_{k,j})$ converges to 0 when $I$ is large, which we show now. Consider $k$ to be fixed. Because each $B_{k,j}$ is independently and identically distributed, if we can show that with high probability, $B_{k,j}$ does not deviate too much from $\mathbb{E}[B_{k,j}$, then we can justify our approximation. The $e^{\varepsilon_{i,j}}$ are independently and identically lognormally distributed. For each $j$, $B_{k,j}$ is the sum of independent random variables. If we assume $N_i \alpha_i$ is bounded and that we have many samples $I$, then Chebyshev's inequality implies that for any $\delta > 0$, as $I \to \infty$,

$$P(|B_{k,j} - \mathbb{E}[B_{k,j}]| > \delta) \to 0 \tag{10}$$

A quantitative bound to (10) for finite $I$ can be derived by Chebyshev's inequality (See "Quantitative bound for deviation of sum of independent random variables"). As such, we are justified in making the approximation that $B_{k,j} \approx \beta_k \beta_0 \equiv \mathbb{E}[B_{k,j}]$ for all $k, j$.

Consequently, if we define $W_k = \beta_k \beta_0$,

$$S_j|\gamma_j \sim \text{Poisson}\left(I\bar{N}e^{\gamma_j}\sum_{k=1}^{K}\mu_{k,j}W_k\right), \quad \gamma_j \sim \text{Normal}(0,\sigma_\gamma^2). \tag{11}$$

We can estimate $W_k$ by viewing this as an (unconstrained) Poisson-lognormal mixture model. Our procedure to calculate the MLE of $W$ and $\sigma_\gamma$ ($\hat{W}$ and $\hat{\sigma_\gamma}$) is the same procedure used to fit RCTD, which is described later (See fitting model (15)). Because the bulk Poisson mean is large for most genes, the Poisson sampling is mostly negligible. Accordingly, with high probability:

$$S_j \approx e^{\gamma_i}\sum_{k=1}^{K}\mu_{k,j}W_k \implies \gamma_j \mid \hat{W} \approx \log(\bar{S}_j) - \log\left(\sum_{k=1}^{K}\mu_{k,j}\hat{W}_k\right) \equiv \hat{\gamma}_j \tag{12}$$

This is an approximation, given that there will be some deviation of $\hat{W}$ from the true $W$. We call this a normalization, because the effect of $\hat{\gamma}_j$ is to re-normalize the cell type means as:

$$\bar{\mu}_{k,j} = \mu_{k,j}e^{\hat{\gamma}_j} \tag{13}$$

**Sequential quadratic programming for MLE estimation of RCTD**

After estimating platform effects, we use RCTD to independently fit each spot as a weighted sum of cell types. Note that since $\gamma_j$ has been replaced with a fixed estimate $\hat{\gamma}_j$, there are no shared parameters across different pixels $i$, except $\sigma_\varepsilon$. As such, we estimate $\beta_{k,i}$ independently for each pixel, given $\sigma_\varepsilon$. The parameter $\sigma_\varepsilon$ is estimated by maximum likelihood via stochastic gradient descent (see "Estimating $\sigma$"). The parameter $\alpha_i$ effectively allows us to re-scale $\beta_i$, so we define $w_{k,i} = \beta_{k,i}e^{\alpha_i}$, which will not be constrained to sum to 1. Next, define:

$$\bar{\lambda}_{i,j}(w_i) = N_i\sum_{k=1}^{K}w_{k,i}\mu_{k,j}e^{\hat{\gamma}_j} = N_i\sum_{k=1}^{K}w_{k,i}\bar{\mu}_{k,j} \tag{14}$$

We will refer to this as the predicted mean of gene $j$ in pixel $i$. The final model is a Poisson-lognormal mixture model:

$$Y_{i,j} \mid \bar{\lambda}_{i,j} \sim \text{Poisson}(e^{\varepsilon_{i,j}}\bar{\lambda}_{i,j}(w_i)), \quad \varepsilon_{i,j} \sim \text{Normal}(0,\sigma_\varepsilon^2) \tag{15}$$

We estimate $w_i^* \geq 0$ as the solution that maximizes the log-likelihood $\mathcal{L}(w_i)$:

$$\max \mathcal{L}(w_i) = \sum_{j=1}^{J}\log P(Y_{i,j} \mid \bar{\lambda}_{i,j}(w_i)) \quad \text{subject to: } w \geq 0 \tag{16}$$

Since the log-likelihood is non-convex, we implement sequential quadratic programming to optimize quadratic approximations and iterate until convergence - typically less than 15 iterations. Since this is the same model as equation (11) in Platform Effect Normalization, we use the same optimization procedure for both models.

Here, we provide the Sequential quadratic programming optimization procedure to optimize the log-likelihood of RCTD (equation (16)). From now on, we consider a fixed pixel $i$ and suppress the notation of $i$. We can directly compute $P(Y_j \mid \bar{\lambda}_j)$ by integrating over the random effect $\varepsilon_j$:

$$\begin{aligned} P(Y_j \mid \bar{\lambda}_j) &= \int_{-\infty}^{\infty} p_\sigma(z)P(Y_j \mid \lambda_j = \bar{\lambda}_j e^z)\mathrm{dz} \\ &= \int_{-\infty}^{\infty} p_\sigma(z)e^{-\bar{\lambda}_j e^z}\frac{(e^z\bar{\lambda}_j)^{Y_j}}{Y_j!}\mathrm{dz} \\ &= Q_{Y_j}(\bar{\lambda}_i) \end{aligned} \tag{17}$$

Here, $p_\sigma$ is the probability density function of $\varepsilon$. From now on, for notational convenience, we will use $\lambda_j$ to denote $\bar{\lambda}_j$. We have defined for $\ell \in \mathbb{Z}^+ \cup \{0\}$:

$$Q_\ell(\lambda) \equiv \int_{-\infty}^{\infty} p_\sigma(z) e^{-\lambda e^z} \frac{(e^z \lambda)^\ell}{\ell!} \mathrm{dz} \tag{18}$$

We will estimate $w^* \geq 0$ as that which maximizes the log-likelihood $\mathcal{L}(w)$:

$$\max_w \mathcal{L}(w) = \sum_{j=1}^{J} \log P(Y_j \mid \lambda_j(w)) = \sum_{j=1}^{J} \log Q_{Y_j}(\lambda_j(w)) \tag{19}$$

Our log likelihood is non-convex. To optimize it, we will apply Sequential Quadratic Programming, an iterative procedure that will be repeated until convergence. Let $w_0$ be the value of $w$ at a given iteration, and let the gradient of $-\mathcal{L}$ be $b(w)$ and the Hessian of $-\mathcal{L}$ be $A(w)$. Then, we can make the following quadratic Taylor approximation to $\mathcal{L}$:

$$-\mathcal{L}(w) \approx -\mathcal{L}(w_0) + b(w_0)^T (w - w_0) + \frac{1}{2}(w - w_0)^T A(w_0)(w - w_0) \tag{20}$$

If we let $d = w - w_0$, then we get the following optimization problem to minimize our approximation of $-\mathcal{L}(w)$.

$$\begin{aligned} \min_d \quad & b(w_0)^T d + \frac{1}{2} d^T A(w_0) d \\ \text{s.t.} \quad & d + w_0 \geq 0 \end{aligned} \tag{21}$$

This is a constrained Quadratic Program (QP). There is one issue we must discuss. This QP will not be well behaved if the Hessian $A(w_0)$ is not positive semi-definite, which can occur due to the non-convexity of $-\mathcal{L}(w)$. As such, we employ an approximation which has been shown to be very effective in non-convex optimization [1]: we set $A(w_0)$ to be the positive semi-definite part of the hessian $H$ of $-\mathcal{L}(w_0)$. Specifically, suppose we have an eigen-decomposition of $H$ as:

$$H = VDV^T \tag{22}$$

Here, $D$ is diagonal matrix of eigenvalues. We obtain the positive semi-definite part of H by taking $D^+ = \max(D, 0)$ and:

$$A = VD^+V^T \tag{23}$$

Effectively, taking the positive semi-definite part of the Hessian will preserve the gradient information and keep the convex part of the quadratic Taylor approximation.

Given a solution of $d^*$, we set

$$w_0 + \alpha * d^* \to w_0 \tag{24}$$

Here, $\alpha$ is the step size, which we have set to 0.3. We repeat this process until convergence. This gives us a solution $w^*$ minimizing $-\mathcal{L}(w)$.

Next, recalling that $\lambda_j(w) = N w^T \bar{\mu}_j$, we will derive an expression for the gradient and hessian of $-\mathcal{L}(w)$:

$$\begin{aligned} b(w) = -\nabla L(w) &= -\sum_{j=1}^{J} \nabla \log Q_{Y_j}(\lambda_j(w)) \\ &= -\sum_{j=1}^{J} \frac{Q'_{Y_j}(\lambda_j(w))}{Q_{Y_j}(\lambda_j(w))} N \bar{\mu}_j \end{aligned} \tag{25}$$

Next, the Hessian:

$$
\begin{aligned}
A(w) = \text{Hess}(-L(w)) &= -\sum_{j=1}^{J} \nabla(\frac{Q'_{Y_j}(\lambda_i(w))}{Q_{Y_j}(\lambda_j(w))}) \cdot N\bar{\mu}_j \\
&= -\sum_{j=1}^{J}(\frac{Q''_{Y_j}(\lambda_i(w))}{Q_{Y_j}(\lambda_j(w))} - (\frac{Q'_{Y_i}(\lambda_i(w))}{Q_{Y_j}(\lambda_j(w))})^2) \cdot (N\bar{\mu}_j)(N\bar{\mu}_j)^T
\end{aligned}
\tag{26}
$$

Consequently, computation of the gradient and Hessian will depend on computing the first and second derivatives of $Q_\ell(\lambda)$. Notice that:

$$
\begin{aligned}
Q'_\ell(\lambda) &\equiv \int_{-\infty}^{\infty} p_\sigma(z) \frac{\partial}{\partial\lambda}(e^{-\lambda_i e^z} \frac{(e^z\lambda_i)^\ell}{\ell!})\text{dz} \\
&= \int_{-\infty}^{\infty} p_\sigma(z) e^{-\lambda_i e^z} \frac{(e^z\lambda_i)^\ell}{\ell!}(-e^z + \frac{\ell}{\lambda})\text{dz} \\
&= \frac{1}{\lambda}(-(\ell+1)Q_{\ell+1}(\lambda) + \ell Q_\ell(\lambda))
\end{aligned}
\tag{27}
$$

As such, we are able to express the derivatives of $Q_\ell$ in terms of $Q_\ell$ and $Q_{\ell+1}$! Likewise, we can compute:

$$
\begin{aligned}
Q''_\ell(\lambda) &= \frac{1}{\lambda}(-(\ell+1)Q'_{\ell+1}(\lambda) + \ell Q'_\ell(\lambda)) - \frac{1}{\lambda^2}(-(\ell+1)Q_{\ell+1}(\lambda) + \ell Q_\ell(\lambda)) \\
&= \frac{1}{\lambda^2}((\ell+1)(\ell+2)Q_{\ell+2}(\lambda) - 2\ell(\ell+1)Q_{\ell+1}(\lambda) + \ell(\ell-1)Q_\ell(\lambda))
\end{aligned}
\tag{28}
$$

Therefore, we have reduced the problems of computing the gradient and Hessian to the problem of computing $Q_\ell(\lambda)$ for all $\ell$ and $\lambda$. For each $\ell$, we create a grid of potential $\lambda$ values and compute $Q_\ell(\lambda)$. We store these results in a matrix, and for new $\lambda$ values, we linearly interpolate to approximate $Q_\ell(\lambda)$.

##### Estimating $\sigma$

Finally, we address the issue of choosing $\sigma$ (for both $\sigma_\gamma$ and $\sigma_\varepsilon$) with the following procedure:

1. Initially set $\sigma = 1$.
2. Estimate the weights $w$ for 500 pixels.
3. Compute the MLE $\sigma^*$ of $\sigma$ for these 500 samples given these values for $w$.

We note two differences in the estimation of $\sigma_\gamma$ for the Platform Effect Normalization: there is only 1 bulk sample, so this is used, rather than 500 pixel, and more iterations over $\alpha_n$ are used below. The MLE for $\sigma$ is found using stochastic gradient descent. We select $\sigma$ to maximize the likelihood across all spots (with $I$ samples):

$$
\mathcal{L}(\sigma, w) = \sum_{i=1}^{I}\sum_{j=1}^{J} \log P(Y_{i,j} \mid \lambda_{i,j}) = \sum_{i=1}^{I}\sum_{j=1}^{J} \log Q_{Y_{i,j}}(\lambda_{i,j}) = \sum_{i,j} \log Q_{Y_{i,j}}(\lambda_{i,j})
\tag{29}
$$

We alternatively maximizing $\mathcal{L}$ with respect to each of $\sigma$ and $w$, as described above. This is guaranteed to converge to a local minimum. In practice, the choice of $\sigma$ does not affect the maximum likelihood weights $w$ too much, so we typically see convergence of $\sigma$ in a couple iterations. The stochastic gradient descent algorithm proceeds as follows:

1. Set $\sigma = \sigma_0$ initial value.
2. Set $\alpha_n = \frac{.0001}{n}$, and repeat the following for $1 \le n \le N_{\text{iter}}$:

3. For each $1 \leq i \leq I, \quad 1 \leq j \leq J$, update:

$$\sigma \leftarrow \sigma + \alpha_n \frac{\partial}{\partial \sigma} \log Q_{Y_{i,j}}(\lambda_{i,j}) \tag{30}$$

The choice of $\alpha_n$ is to guarantee convergence to the global minimum since $\sum_n^\infty \alpha_n = \infty$, but $\sum_n^\infty \alpha_n^2 < \infty$. Traces of $\sigma$ throughout this procedure imply convergence (Supplementary Fig. 10). A typical choice for $N_{\text{iter}}$ is 10. One remaining detail for this procedure is the calculation of $\frac{\partial}{\partial \sigma} \log Q_{Y_{i,j}}(\lambda_{i,j})$:

$$\begin{aligned}\frac{\partial}{\partial \sigma} \log Q_{Y_{i,j}}(\lambda_{i,j}) &= \frac{1}{Q_{Y_{i,j}}(\lambda_{i,j})} \frac{\partial}{\partial \sigma} Q_{Y_{i,j}}(\lambda_{i,j}) \\ &= \frac{1}{Q_{Y_{i,j}}(\lambda_{i,j})} \int_{-\infty}^{\infty} \frac{\partial p_\sigma(z)}{\partial \sigma} e^{-\lambda_{i,j} e^z} \frac{(e^z \lambda_{i,j})^{Y_{i,j}}}{(Y_{i,j})!} \mathrm{dz}\end{aligned} \tag{31}$$

Finally, we must calculate $\frac{\partial p_\sigma(z)}{\partial \sigma}$. We define:

$$p_\sigma(z) = \begin{cases} \frac{C}{\sqrt{2\pi}\sigma} e^{-\frac{z^2}{2\sigma^2}}, & |z| \leq 3\sigma \\ \frac{Ca}{\sigma(z/\sigma - c)^2}, & |z| > 3\sigma \end{cases} \tag{32}$$

Here, $p_\sigma$ is a Normal Distribution, modified to be heavy tailed. The intuition here is that we want to allow for the (inevitable) possibility of outliers. Inverse square tails are almost as heavy tailed as possible for probability distributions. We choose $c$ and $a$ so that $p_\sigma$ is continuously differentiable at the boundary $|z| = 3\sigma$. This results in $c = 7/3$ and $a = \frac{4}{9} e^{-9/2}/\sqrt{2\pi}$. $C$ is a normalizing constant which is chosen to make $p_\sigma$ integrate to 1. Note that these choices of constants hold for all $\sigma$. It follows that:

$$\frac{\partial p_\sigma(z)}{\partial \sigma} = \frac{-1}{\sigma} p_\sigma(z) + \begin{cases} \frac{z^2}{\sigma^3} p_\sigma(z) & |z| \leq 3\sigma \\ \frac{-2/\sigma^2}{z/\sigma - c} p_\sigma(z) & |z| > 3\sigma \end{cases} \tag{33}$$

**Quantitative bound for deviation of sum of independent random variables**

Here, we provide a quantitative bound for equation (10). Recall:

$$B_{k,j} = \frac{1}{I} \sum_{i=1}^{I} \frac{N_i}{\bar{N}} \beta_{k,i} \exp(\alpha_i + \varepsilon_{i,j}) \tag{34}$$

Recall we have independently lognormally distributed $e^{\varepsilon_{i,j}}$. For each $j$, $B_{k,j}$ is the sum of independent random variables. If we assume $N_i \alpha_i$ is bounded and that we have many samples $I$, then Chebyshev's inequality implies that for any $\delta > 0$:

$$\begin{aligned}P(|B_{k,j} - \mathbb{E}[B_{k,j}]| > \delta) &\leq \frac{\mathrm{Var}(B_{k,j})}{\delta^2} \\ &= \frac{1}{(I\bar{N}\delta)^2} \sum_{i=1}^{I} (N_i \beta_{k,i} e^{\alpha_i})^2 \mathrm{Var}(e^{\varepsilon_{i,j}}) \\ &\leq \frac{\max_{1 \leq i \leq I} (N_i e^{\alpha_i})^2 (e^{\sigma_\varepsilon^2}(e^{\sigma_\varepsilon^2} - 1))}{I \delta^2 \bar{N}^2},\end{aligned} \tag{35}$$

where we have used the variance of a lognormal distribution. Therefore, with high probability, $\beta_{k,j}$ will not deviate far from its mean for large $I$.

**Expected cell type-specific gene expression**

Once $\beta$ has been estimated, we probabilistically assign reads to cell types. This is accomplished by our model of sampling gene expression: for each pixel $1 \le i \le I$ and for each read $1 \le n \le N_i$, we select cell type $1 \le \theta_{n,i} \le K$ and gene $1 \le z_{n,i} \le J$ as:

$$P(\theta_{n,i} = k \mid \beta) = \beta_{k,i}, \quad P(z_{n,i} = j \mid \theta_{n,i}, \gamma, \varepsilon) = \delta_{i,\theta_{n,i},j} \tag{36}$$

Here, we define:

$$\log(\delta_{i,k,j}) = \alpha_i + \log(\hat{\mu}_{k,j}) + \gamma_j + \varepsilon_{i,j} \tag{37}$$

We can use Bayes' Theorem to calculate the probability that each read $n$ of gene $i$ came from cell type $j$ in sample $k$. We assume that the reads are conditionally independent given $\beta, \gamma, \varepsilon$:

$$\begin{aligned} P(\theta_{n,i} = k \mid \beta, z_{n,i} = j, \gamma, \varepsilon) &\propto P(\theta_{n,i} = k, z_{n,i} = j \mid \beta, \gamma, \varepsilon) \\ &= P(\theta_{n,i} = k \mid \beta) P(z_{n,i} = j \mid \beta, \theta_{n,i} = k, \gamma, \varepsilon) \\ &= \beta_{k,i}\delta_{i,k,j} \end{aligned} \tag{38}$$

This implies that:

$$P(\theta_{n,i} = k \mid \beta, z_{n,i} = j, \gamma, \varepsilon) = \frac{\beta_{k,i}\delta_{i,k,j}}{\sum_{k'=1}^{K} \beta_{k',i}\delta_{i,k',j}} = \frac{\beta_{k,i}\hat{\mu}_{k,j}}{\sum_{k'=1}^{K} \beta_{k',i}\hat{\mu}_{k',j}} \tag{39}$$

This conditional probability does not depend on $\gamma, \varepsilon$, since the ratio (across $k$) of $\delta_{i,k,j}$ is fixed. This implies (by tower law):

$$P(\theta_{n,i} = k \mid \beta, z_{n,i} = j) = \mathbb{E}_{\gamma,\varepsilon}[P(\theta_{n,i} = k \mid \beta, z_{n,i} = j, \gamma, \varepsilon)] = \frac{\beta_{k,i}\hat{\mu}_{k,j}}{\sum_{k'=1}^{K} \beta_{k',i}\hat{\mu}_{k',j}} \tag{40}$$

Therefore, we can use Equation (40) to calculate the probability that each UMI came from each cell type given $\beta_{k,i}$. Intuitively, this probability is a competition between different cell types and is proportional to (1) the proportion of the cell type on a spot and (2) the probability of observing the gene in the cell type for a single UMI. Consequentially, conditional on the observed gene counts, we can calculate in pixel $i$ the expectation of gene $j$ originating from each cell type $k$, $Y_{i,k,j}$, as proportional to the observed gene counts $Y_{i,j}$:

$$\mathbb{E}[Y_{i,k,j} \mid \beta, Y_{i,j}] = \mathbb{E}[\sum_{i=1}^{I} \mathbb{I}[\theta_{n,i} = k, z_{n,i} = j] \mid \beta, Y_{i,k}] = \frac{Y_{i,j}\beta_{k,i}\hat{\mu}_{k,j}}{\sum_{k'=1}^{K} \beta_{k',i}\hat{\mu}_{k',j}} \tag{41}$$

**Supplementary Table 1: Estimated Parameters**

| Target Dataset | $\sigma_\gamma$ | $\sigma_\varepsilon$ |
|---|---|---|
| snucRNA-seq Cerebellum | 0 | 0.77 |
| scRNA-seq Cerebellum | 1.28 | 0.88 |
| Slide-seq Cerebellum | 0.94 | 0.95 |
| Slide-seq Hippocampus | 0.51 | 0.80 |
| Visium Hippocampus | 0.70 | 0.45 |

**Equations for schematic**

$$\begin{aligned} Y_{i,j} \mid \lambda_{i,j} &\sim \text{Poisson}(N_i \lambda_{i,j}) \\ \lambda_{i,j} \mid \beta, \gamma, \mu &\sim \mathcal{LN}(e^{\gamma_j + \alpha_i} \sum_{k=1}^{K} \beta_{i,k}\mu_{k,j}) \end{aligned} \tag{42}$$

### Supplementary Experimental Methods

##### Animal Handling

All procedures involving animals at the Broad Institute were conducted in accordance with the US National Institutes of Health Guide for the Care and Use of Laboratory Animals under protocol number 0120-09-16.

##### Transcardial Perfusion

C57BL/6J mice were anesthetized by administration of isoflurane in a gas chamber flowing 3% isoflurane for 1 minute. Anesthesia was confirmed by checking for a negative tail pinch response. Animals were moved to a dissection tray and anesthesia was prolonged via a nose cone flowing 3% isoflurane for the duration of the procedure. Transcardial perfusions were performed with ice cold pH 7.4 HEPES buffer containing 110 mM NaCl, 10 mM HEPES, 25 mM glucose, 75 mM sucrose, 7.5 mM MgCl2, and 2.5 mM KCl to remove blood from brain and other organs sampled. The appropriate organs were removed and frozen for 3 minutes in liquid nitrogen vapor and moved to -80C for long term storage.

##### Tissue Handling

Fresh frozen tissue was warmed to -20 C in a cryostat (Leica CM3050S) for 20 minutes prior to handling. Tissue was then mounted onto a cutting block with OCT and sliced at a 5° cutting angle at 10 µm thickness. Pucks were then placed on the cutting stage and tissue was maneuvered onto the pucks. The tissue was then melted onto the puck by moving the puck off the stage and placing a finger on the bottom side of the glass. The puck was then removed from the cryostat and placed into a 1.5 mL eppendorf tube. The sample library was then prepared as below. The remaining tissue was re-deposited at -80 C and stored for processing at a later date.

##### Puck preparation and sequencing

Pucks were prepared as described recently using barcoded beads synthesized in-house on an Akta Oligopilot 10 according to the updated Slide-seqV2 protocol [2]. Pucks were sequenced using a monobase-encoding sequencing-by-ligation approach also described in the updated protocol. We used slide-seq tools for alignment and processing of Slide-seq data.

Pucks were generated using one of two separate bead batches with the oligo sequences listed below:

Batch 1:

5'-
TTT_PC_GCCGGTAATACGACTCACTATAGGGCTACACGACGCTCTTCCGATCTJJJJJJJJTCTTCAGCGTTCCCGAGAJ
JJJJJJTCNNNNNNNNT25

Batch 2:

5'-
TTT_PC_GCCGGTAATACGACTCACTATAGGGCTACACGACGCTCTTCCGATCTJJJJJJJJTCTTCAGCGTTCCCGAGAJ
JJJJJNNNNNNNNVVT30

"PC" designates a photocleavable linker; "J" represents bases generated by split-pool barcoding, such that every oligo on a given bead has the same J bases; "N" represents bases generated by mixing, so every oligo on a given bead has different N bases; and "TX" represents a sequence of X thymidines. "V" represents bases which may contain A, C, G but not T.

**Slide-seqV2 library preparation**

RNA Hybridization:
Pucks in 1.5 mL tubes were immersed in 200 µL of hybridization buffer (6x SSC with 2 U/µL Lucigen NxGen RNAse inhibitor) for 15 minutes at room temperature to allow for binding of the RNA to the oligos on the beads.

First Strand Synthesis
Subsequently, first strand synthesis was performed by incubating the pucks in RT solution for 30 minutes at room temperature followed by 1.5 hours at 52 °C.

RT solution:
115 µL H2O
40 µL Maxima 5x RT Buffer (Thermofisher, EP0751**)**
20 µL 10 mM dNTPs (NEB N0477L)
5 µL RNase Inhibitor (Lucigen 30281)
10 µL 50 µM Template Switch Oligo **(**Qiagen #339414YCO0076714)
10 µL Maxima H- RTase (Thermofisher, EP0751)

Tissue Digestion:
200 µL of 2x tissue digestion buffer was then added directly to the RT solution and the mixture was incubated at 37 °C for 30 minutes.

2x tissue digestion buffer:
200 mM Tris-Cl pH 8
400 mM NaCl
4% SDS
10 mM EDTA
32 U/mL Proteinase K (NEB P8107S)

Second Strand Synthesis:
The solution was then pipetted up and down vigorously to remove beads from the surface, and the glass substrate was removed from the tube using forceps and discarded. 200 µL of Wash Buffer was then added to the 400 µL of tissue clearing and RT solution mix and the tube was then centrifuged for 2 minutes at 3000 RCF. The supernatant was then removed from the bead pellet, the beads were resuspended in 200 µL of Wash Buffer, and were centrifuged again. This was repeated a total of three

times. The supernatant was then removed from the pellet. The beads were then resuspended in 200 µL of ExoI mix and incubated at 37 °C for 50 minutes.

Wash Buffer:
10 mM Tris pH 8.0
1 mM EDTA
0.01% Tween-20

ExoI mix:
170 µL H20
20 µL ExoI buffer
10 µL ExoI (NEB M0568)

After ExoI treatment the beads were centrifuged for 2 minutes at 3000 RCF. The supernatant was then removed from the bead pellet, the beads were resuspended in 200 µL of Wash Buffer, and were centrifuged again. This was repeated a total of three times. The supernatant was then removed from the pellet. The pellet was then resuspended in 200 µL of 0.1 N NaOH and incubated for 5 minutes at room temperature. To quench the reaction, 200 µL of Wash Buffer was added and beads were centrifuged for 2 minutes at 3000 RCF. The supernatant was then removed from the bead pellet, the beads were resuspended in 200 µL of Wash Buffer, and were centrifuged again. This was repeated a total of three times. Second Strand Synthesis was then performed on the beads by incubating the pellet in 200 µL of Second Strand Mix at 37 °C for 1 hour.

Second Strand Synthesis mix:
133 µL H2O
40 µL Maxima 5x RT Buffer
20 µL 10 mM dNTPs
2 µL 1 mM dN-SMRT oligo
5 µL Klenow Enzyme (NEB M0210)

After Second Strand Synthesis, 200 µL of Wash Buffer was added and the beads were centrifuged for 2 minutes at 3000 RCF. The supernatant was then removed from the bead pellet, the beads were resuspended in 200 µL of Wash Buffer, and were centrifuged again. This was repeated a total of three times.

Library Amplification:
200 µL of water was then added to the bead pellet and the beads were centrifuged for 2 minutes at 3000 RCF. The supernatant was then removed from the bead pellet and the beads were resuspended in 50 µL of library PCR mix and moved into a 200 µL PCR strip tube. PCR was then performed as outlined below:

Library PCR mix:
22 µL H2O
25 µL of Terra Direct PCR mix Buffer (Takara Biosciences 639270)
1 µL of Terra Polymerase (Takara Biosciences 639270)
1 µL of 100 µM Truseq PCR primer (IDT)
1 µL of 100 µM SMART PCR primer (IDT)

PCR program:
95 °C 3 minutes

4 cycles of:
98 °C 20 seconds
65 °C 45 seconds
72 °C 3 minutes

9 cycles of:
98 °C 20 seconds
67 °C 20 seconds
72 °C 3 minutes

Then:
72 °C 5 minutes
Hold at 4 °C

PCR cleanup and Nextera Tagmentation:

Samples were cleaned with Ampure XP (Beckman Coulter A63880) beads in accordance with manufacturer’s instructions at a 0.6x bead/sample ratio (30 µL of beads to 50 µL of sample) and resuspended in 50 µL of water. The cleanup procedure was repeated, this time resuspending in a final volume of 10 µL. 1 µL of the library was quantified on an Agilent Bioanalyzer High sensitivity DNA chip (Agilent 5067-4626). Then, 600 pg of cDNA was taken from the PCR product and prepared into Illumina sequencing libraries through tagmentation using the Nextera XT kit (Illumina FC-131-1096). Tagmentation was performed according to manufacturer's instructions and the library was amplified with primers Truseq5 and N700 series barcoded index primers. The PCR program was as follows:

PCR program:
72 °C for 3 minutes
95 °C for 30 seconds

12 cycles of:
95 °C for 10 seconds
55 °C for 30 seconds
72 °C for 30 seconds

72 °C for 5 minutes
Hold at 4 °C

Samples were cleaned with Ampure XP (Beckman Coulter A63880) beads in accordance with manufacturer’s instructions at a 0.6x bead/sample ratio (30 µL of beads to 50 µL of sample) and resuspended in 10 µL of water. 1 µL of the library was quantified on an Agilent Bioanalyzer High sensitivity DNA chip (Agilent 5067-4626). Finally, the library concentration was normalized to 4 nM for sequencing. Samples were sequenced on the Illumina NovaSeq S2 flowcell 100 cycle kit with 12 samples per run (6 samples per lane) with the read structure 44 bases Read 1, 8 bases i7 index read, 50 bases Read 2. Each puck received approximately 200-400 million reads, corresponding to 3,000-5,000 reads per bead.

#### Supplementary Figures

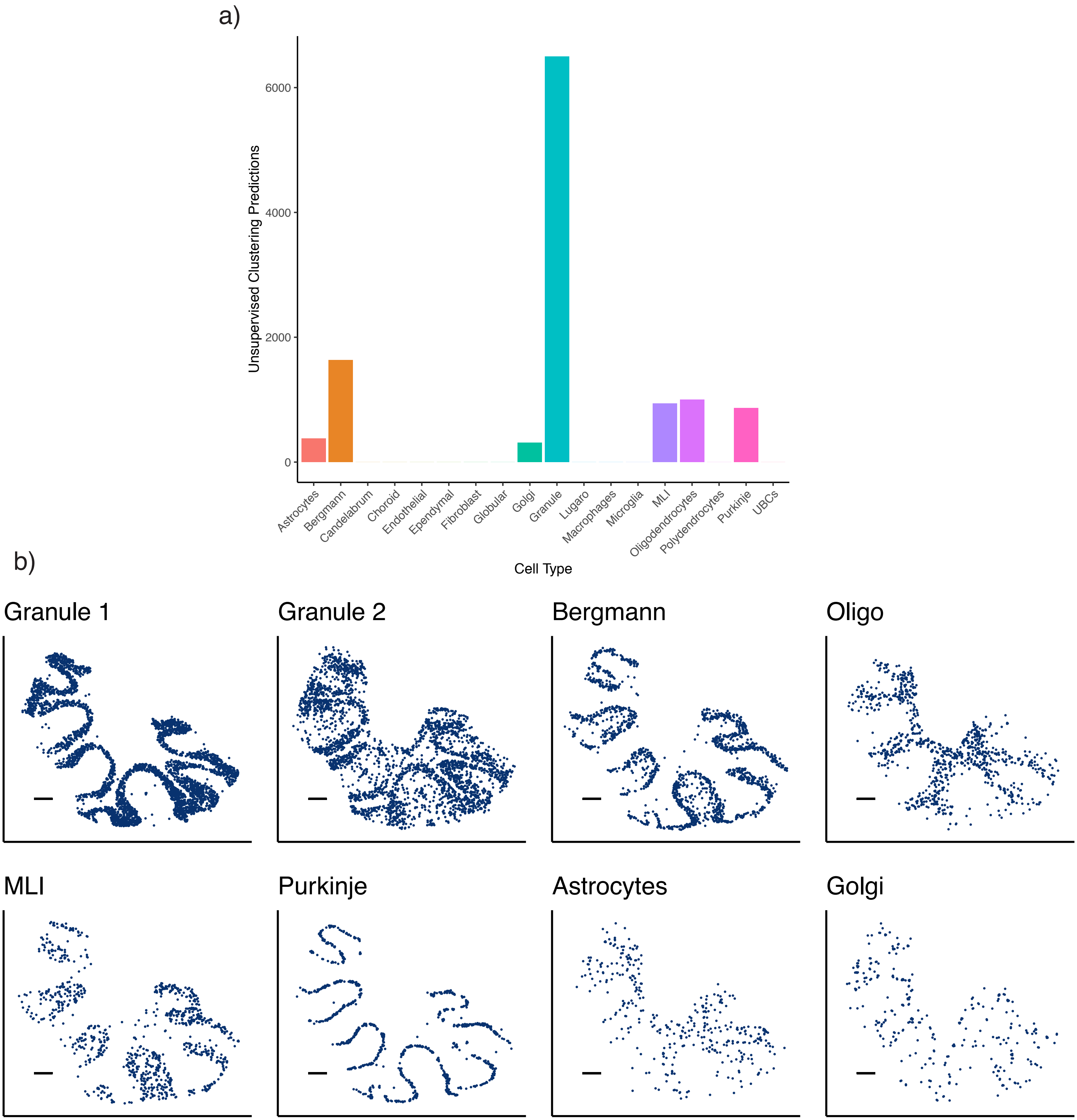
a)
Unsupervised Clustering Predictions
6000
4000
2000
0
Astrocytes
Bergmann
Candelabrum
Choroid
Endothelial
Ependymal
Fibroblast
Globular
Golgi
Granule
Lugaro
Macrophages
Microglia
MLI
Oligodendrocytes
Polydendrocytes
Purkinje
UBCs
Cell Type
b)
Granule 1
Granule 2
Bergmann
Oligo
MLI
Purkinje
Astrocytes
Golgi

Supplementary Figure 1: Predicted spatial localization of cell types by unsupervised clustering in Slide-seq cerebellum.

a) Barplot of counts for predicted pixels assigned to each cell type. Calls obtained unsupervised clustering. Note that there are two granule clusters, and only 1 cluster for both MLI1 and MLI2.

b) Predicted spatial locations of each cell type by unsupervised clustering.

All scale bars 250 microns.

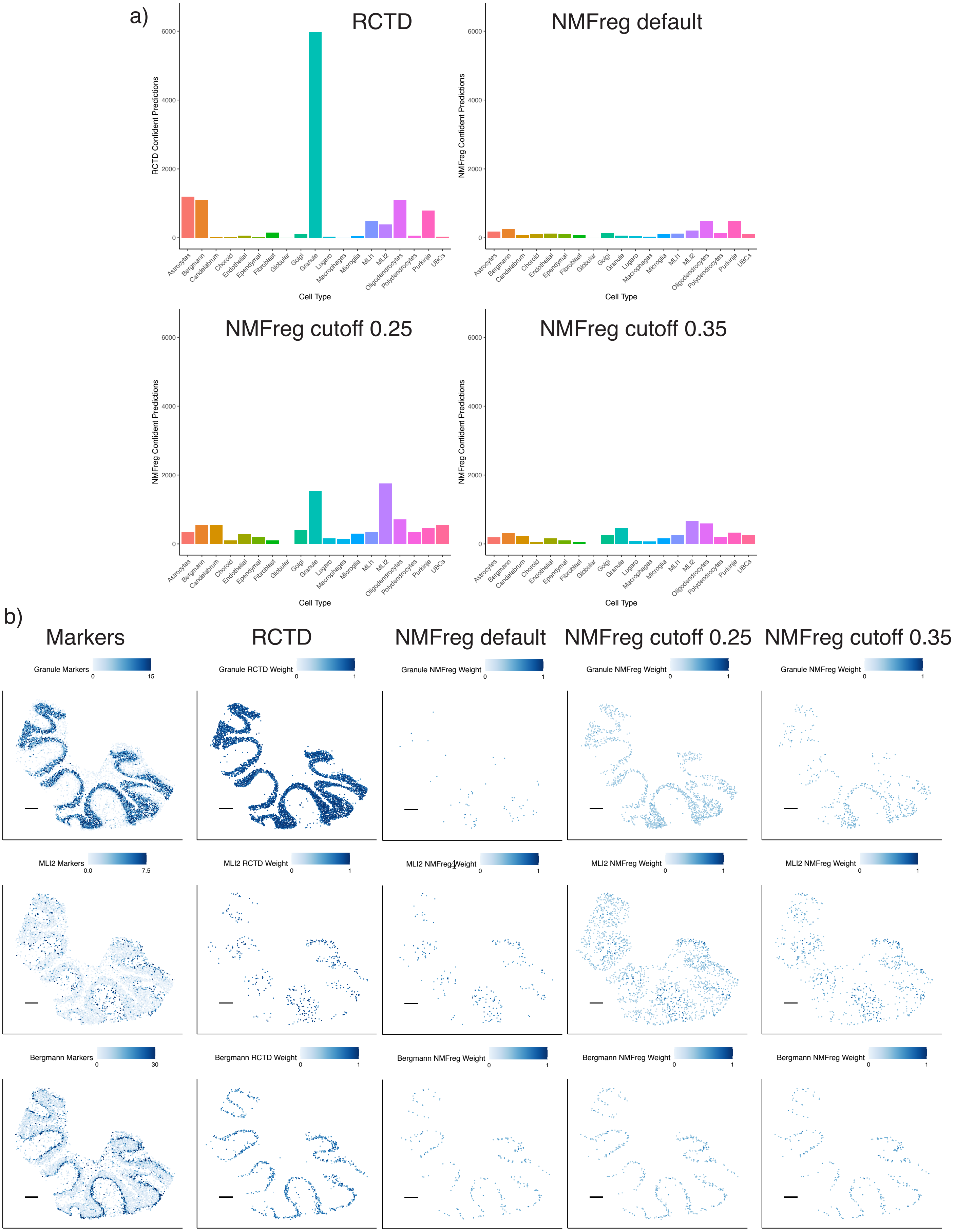

a)
RCTD
NMFreg default
NMFreg cutoff 0.25
NMFreg cutoff 0.35
RCTD Confident Predictions
NMFreg Confident Predictions
Cell Type
Astrocytes
Bergmann
Candelabrum
Choroid
Endothelial
Ependymal
Fibroblast
Globular
Golgi
Granule
Lugaro
Macrophages
Microglia
MLI1
MLI2
Oligodendrocytes
Polydendrocytes
Purkinje
UBCs
b)
Markers
RCTD
NMFreg default
NMFreg cutoff 0.25
NMFreg cutoff 0.35
Granule Markers
Granule RCTD Weight
Granule NMFreg Weight
MLI2 Markers
MLI2 RCTD Weight
MLI2 NMFreg Weight
Bergmann Markers
Bergmann RCTD Weight
Bergmann NMFreg Weight

Supplementary Figure 2: Comparison of NMFreg and RCTD.

a) Barplot of counts for confidently predicted pixels assigned to each cell type. Calls obtained by RCTD (top left) or NMFreg (other three panels). NMFreg confidence was determined either by default (top right), or by a constant proportion cutoff of 0.25 (bottom left) or 0.35 (bottom right).

b) Confidently predicted spatial localization of cell types by RCTD and NMFreg for granule, molecular layer interneurons 2 (MLI2), and Bergmann. Left: expression (counts per 500) (represented by color) of marker genes. Middle: predicted spatial locations of a cell type by RCTD, with color representing predicted cell type proportion. Column 2: predicted spatial locations of a cell type by NMFreg, with color representing predicted cell type proportion. NMFreg underpredicted granule cells in the granular layer and instead incorrectly overpredicted MLI2 in this layer [3]. NMFreg confidence was determined either by default (column 3), or by a constant proportion cutoff of 0.25 (column 4) or 0.35 (column 5).

All scale bars 250 microns.

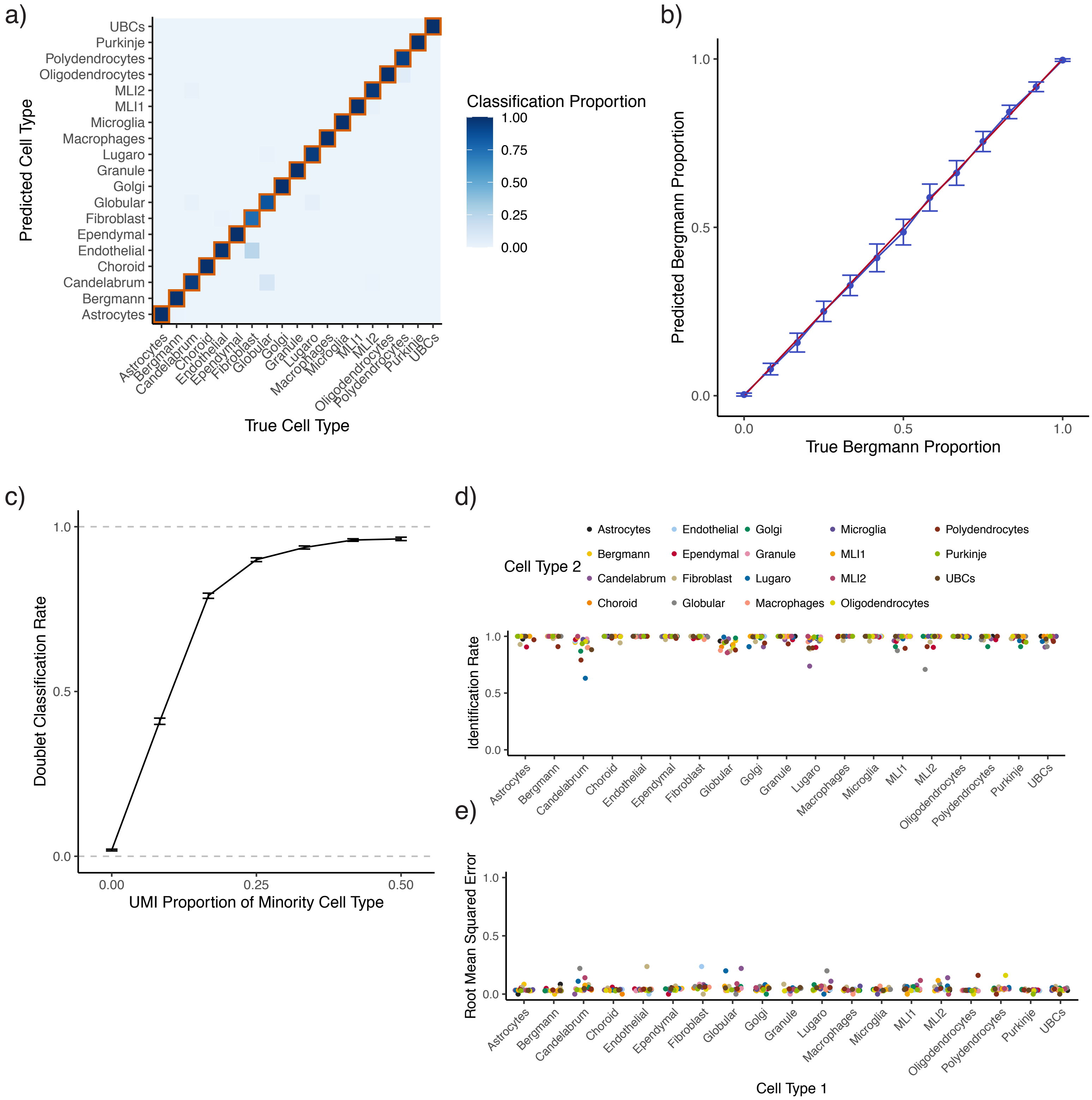
a)
Predicted Cell Type
UBCs
Purkinje
Polydendrocytes
Oligodendrocytes
MLI2
MLI1
Microglia
Macrophages
Lugaro
Granule
Golgi
Globular
Fibroblast
Ependymal
Endothelial
Choroid
Candelabrum
Bergmann
Astrocytes
True Cell Type
Classification Proportion
1.00
0.75
0.50
0.25
0.00
b)
Predicted Bergmann Proportion
True Bergmann Proportion
0.0
0.5
1.0
c)
Doublet Classification Rate
UMI Proportion of Minority Cell Type
0.00
0.25
0.50
d)
Cell Type 2
Identification Rate
e)
Root Mean Squared Error
Cell Type 1

Supplementary Figure 3: Robust Cell Type Decomposition (RCTD) accurately decomposes mixtures of cells.

All: RCTD was trained on the single-nucleus RNA-seq cerebellum dataset and tested on a the same dataset.

a) Confusion matrix for RCTD on within-reference cell type assignment for single cells.

b) Rate of doublet classification of simulated mixtures of single cells, with 95% confidence intervals. The x-axis represents the proportion of UMIs sourced from the minority cell type, ranging from 0% (true singlet) to 50% (full doublet) ($n$ between 5130 and 10260 simulations per condition).

c) Predicted Bergmann proportion as a function of true Bergmann proportion on simulated Bergmann-Purkinje doublets. The red line is the identity line, and the blue line is the average and standard deviation ($n = 30$ simulations per condition) of prediction.

d) Metrics for performance of RCTD on simulated doublets ($n = 390$ simulations per cell type pair). Column represents cell type 1, and color represents cell type 2. Top: RCTD assigns doublets to two cell types. Identification rate is the percentage of confident calls correctly identifying cell class 1.

e) Bottom: RMSE of predicted vs true cell type proportion (as in (c)).

All scale bars 250 microns.

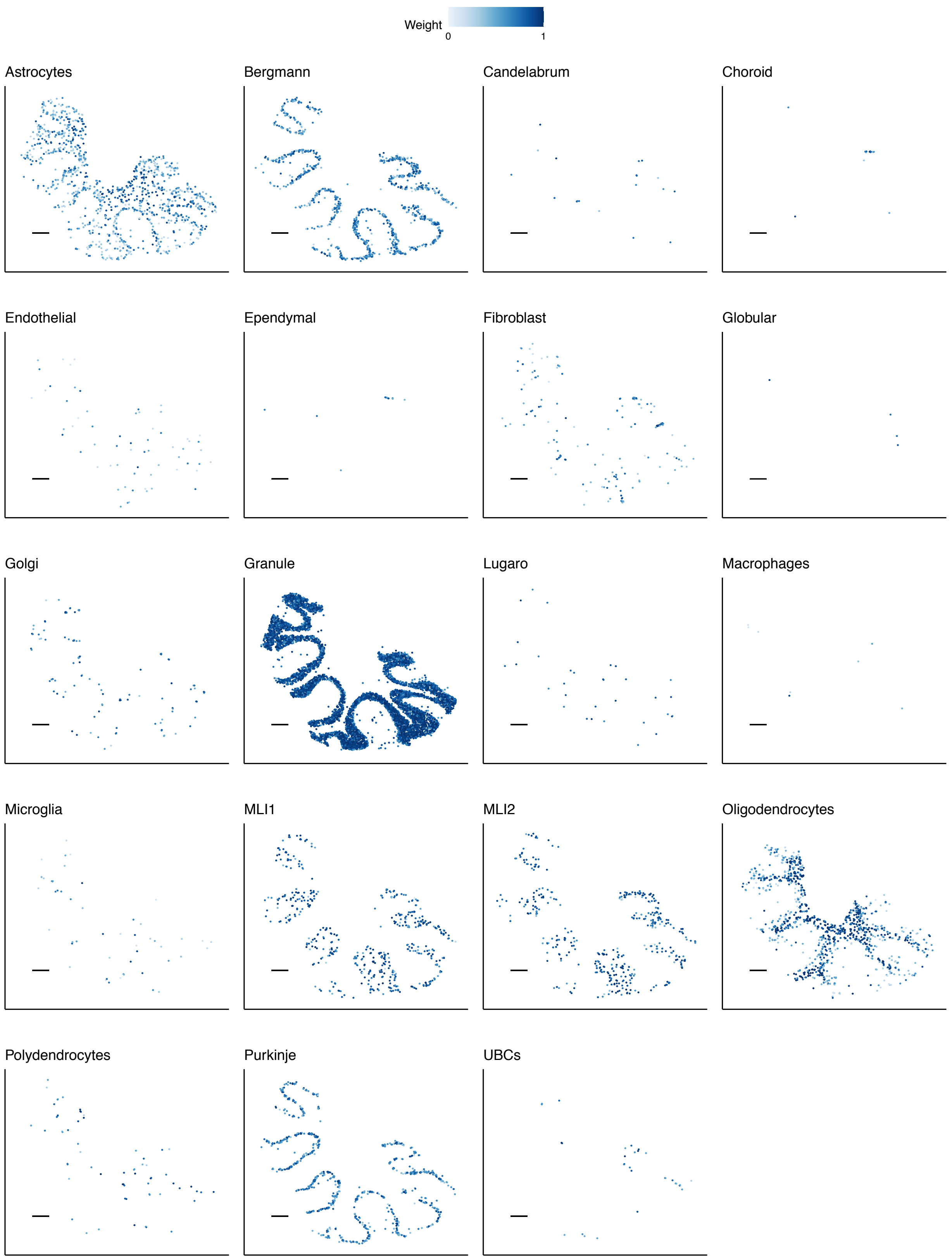
Weight
0
1
Astrocytes
Bergmann
Candelabrum
Choroid
Endothelial
Ependymal
Fibroblast
Globular
Golgi
Granule
Lugaro
Macrophages
Microglia
MLI1
MLI2
Oligodendrocytes
Polydendrocytes
Purkinje
UBCs

Supplementary Figure 4: Predicted spatial localization of cell types by RCTD in Slide-seq cerebellum.

Predicted spatial locations of each cell type, with color representing predicted cell type proportion. All scale bars are 250 microns.

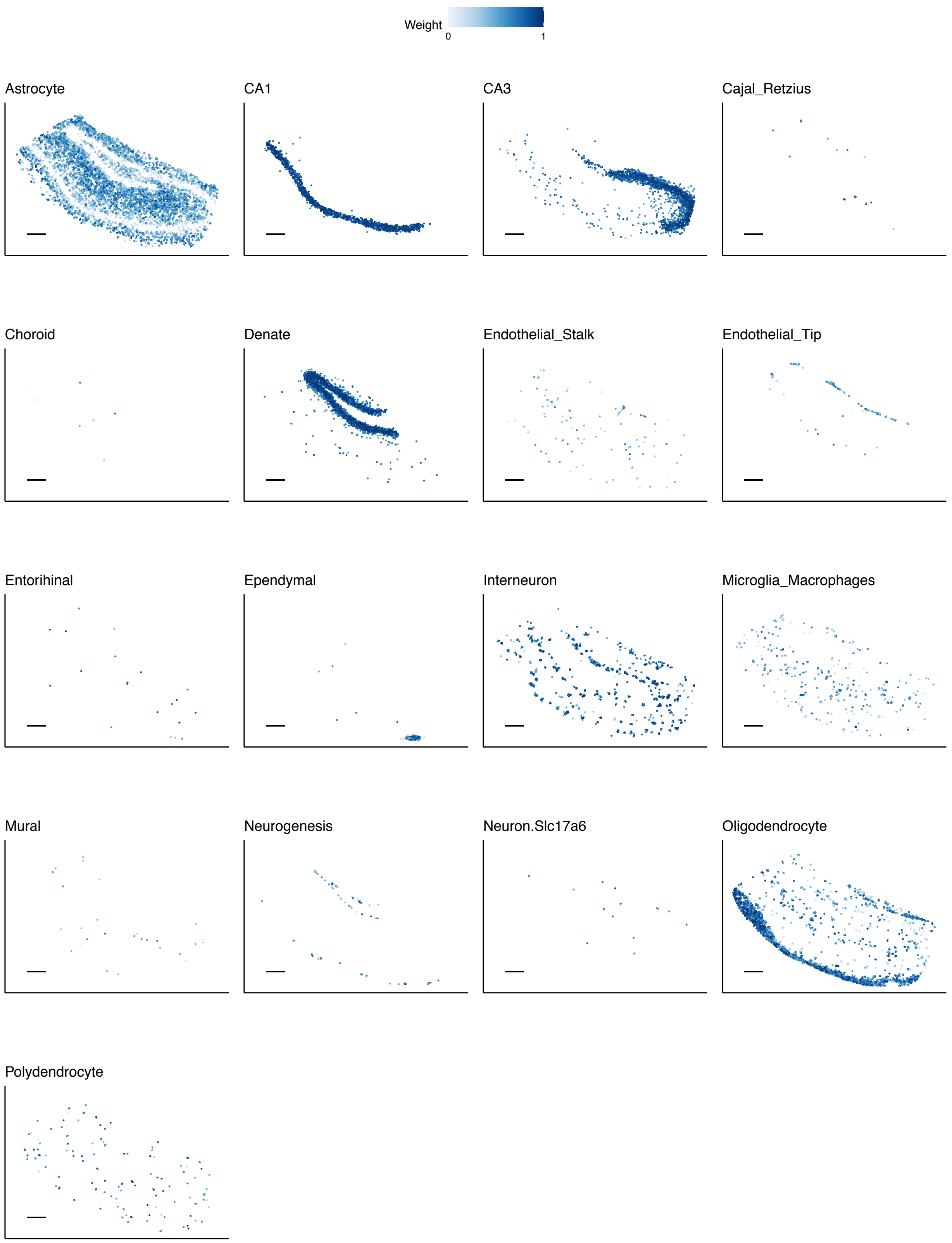
Weight
0
1
Astrocyte
CA1
CA3
Cajal_Retzius
Choroid
Denate
Endothelial_Stalk
Endothelial_Tip
Entorihinal
Ependymal
Interneuron
Microglia_Macrophages
Mural
Neurogenesis
Neuron.Slc17a6
Oligodendrocyte
Polydendrocyte

Supplementary Figure 5: Predicted spatial localization of cell types by RCTD in Slide-seq hippocampus.

Predicted spatial locations of each cell type, with color representing predicted cell type proportion. All scale bars are 250 microns.

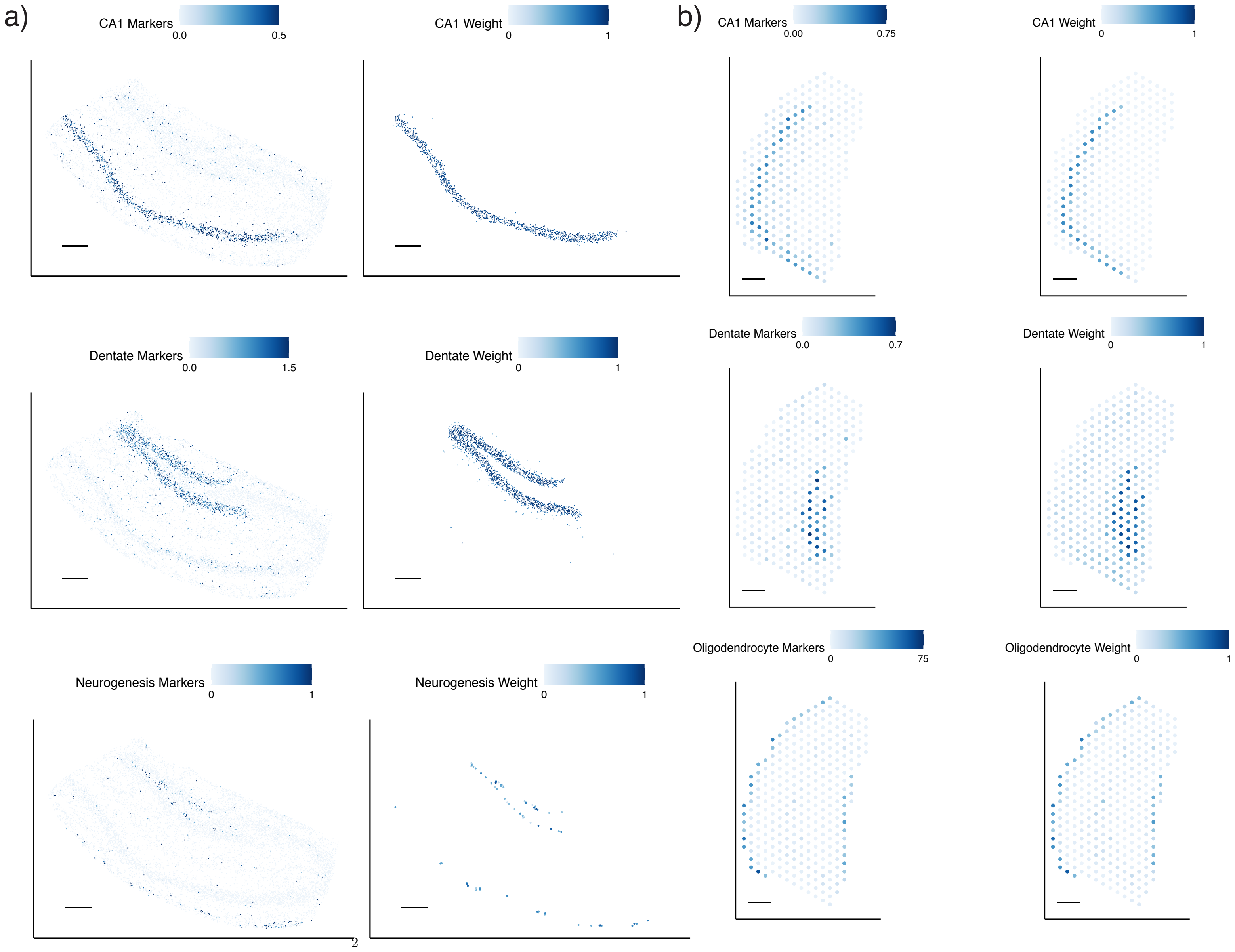
a)
CA1 Markers
0.0
0.5
CA1 Weight
0
1
Dentate Markers
0.0
1.5
Dentate Weight
0
1
Neurogenesis Markers
0
1
Neurogenesis Weight
0
1
2
b)
CA1 Markers
0.00
0.75
CA1 Weight
0
1
Dentate Markers
0.0
0.7
Dentate Weight
0
1
Oligodendrocyte Markers
0
75
Oligodendrocyte Weight
0
1

Supplementary Figure 6: Comparison of marker expression and predicted spatial localization of cell types by RCTD in Slide-seq and Visium hippocampus

a) Slide-seq hippocampus. Left: expression (counts per 500) (represented by color) of marker genes of each cell type. Right: predicted spatial locations of a cell type by RCTD, with color representing predicted cell type proportion.

b) Visium hippocampus. Left: expression (counts per 500) (represented by color) of marker genes of each cell type. Right: predicted spatial locations of a cell type by RCTD, with color representing predicted cell type proportion.

All scale bars 250 microns.

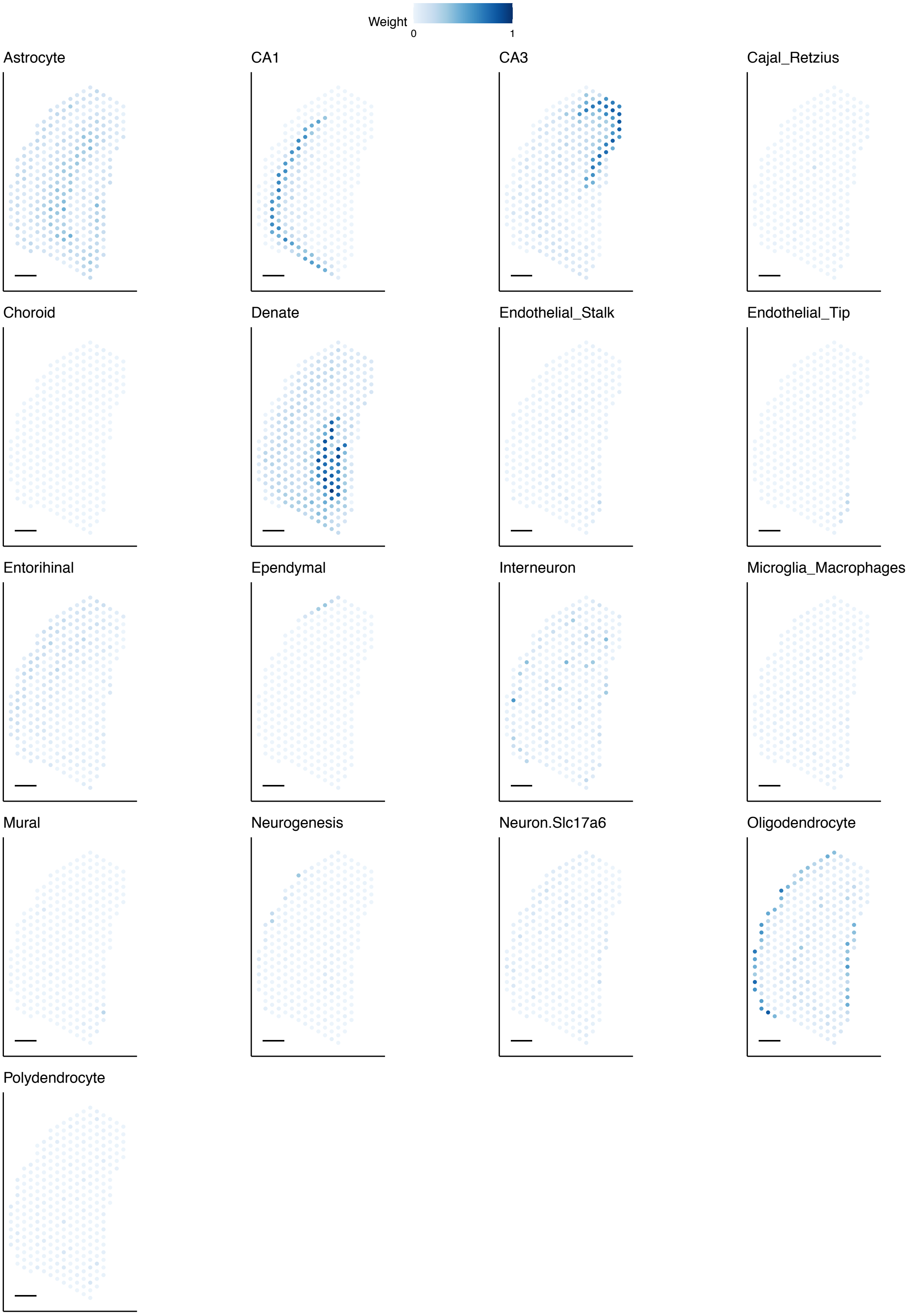
Weight
0
1
Astrocyte
CA1
CA3
Cajal_Retzius
Choroid
Denate
Endothelial_Stalk
Endothelial_Tip
Entorihinal
Ependymal
Interneuron
Microglia_Macrophages
Mural
Neurogenesis
Neuron.Slc17a6
Oligodendrocyte
Polydendrocyte

Supplementary Figure 7: Predicted spatial localization of cell types by RCTD in Visium hippocampus.

Predicted spatial locations of each cell type, with color representing predicted cell type proportion. pixels were not constrained in the number of cell types present. All scale bars are 250 microns.

a)

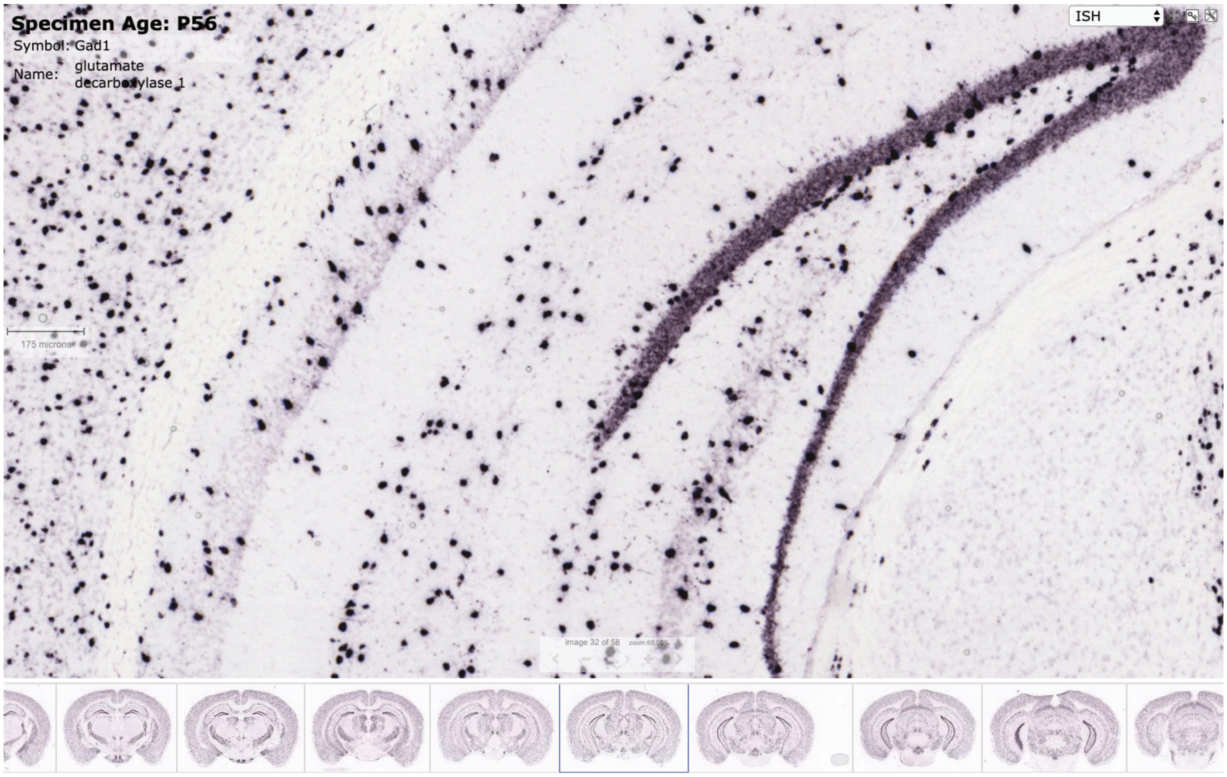
Specimen Age: P56
Symbol: Gad1
Name: glutamate decarboxylase 1
ISH
175 microns

b)

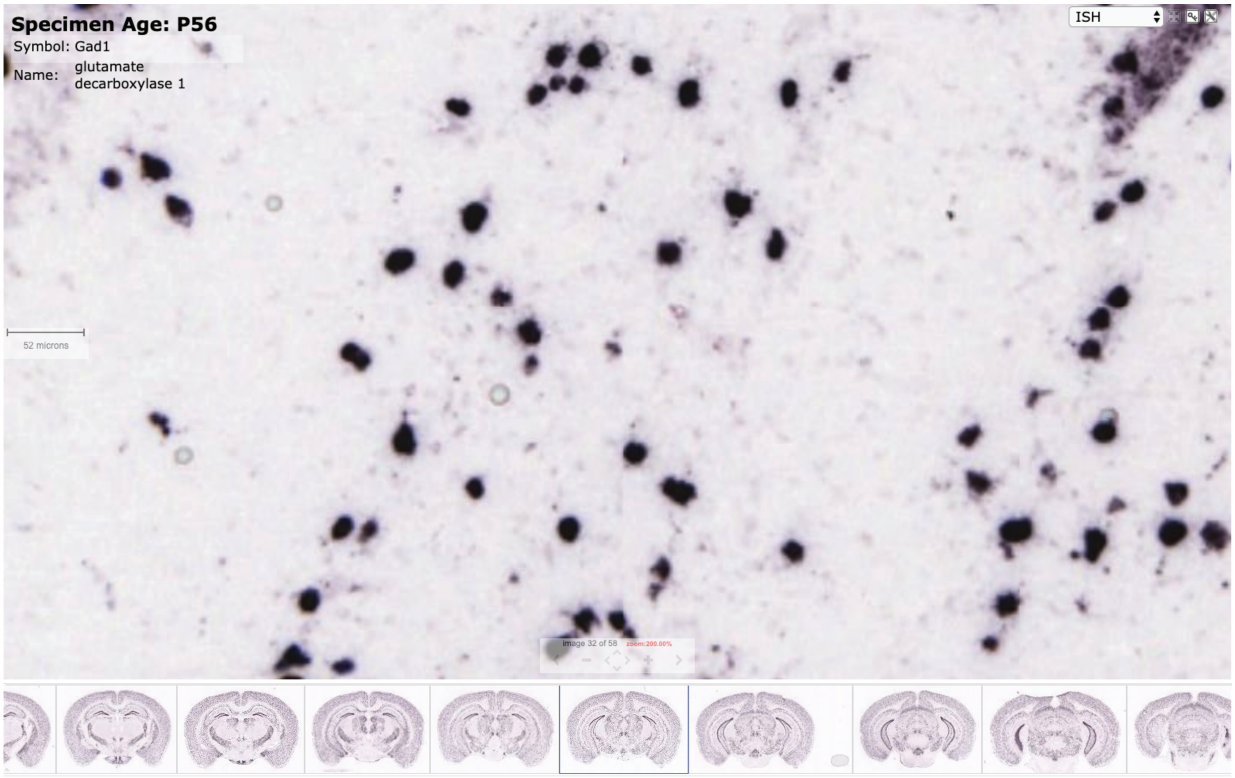
Specimen Age: P56
Symbol: Gad1
Name: glutamate decarboxylase 1
ISH
52 microns

Supplementary Figure 8: In Situ Hybridization (ISH) of *Gad1*, an interneuron marker gene, from the Allen Brain Atlas [4].

a) Scale bar 175 microns.

b) Scale bar 52 microns.

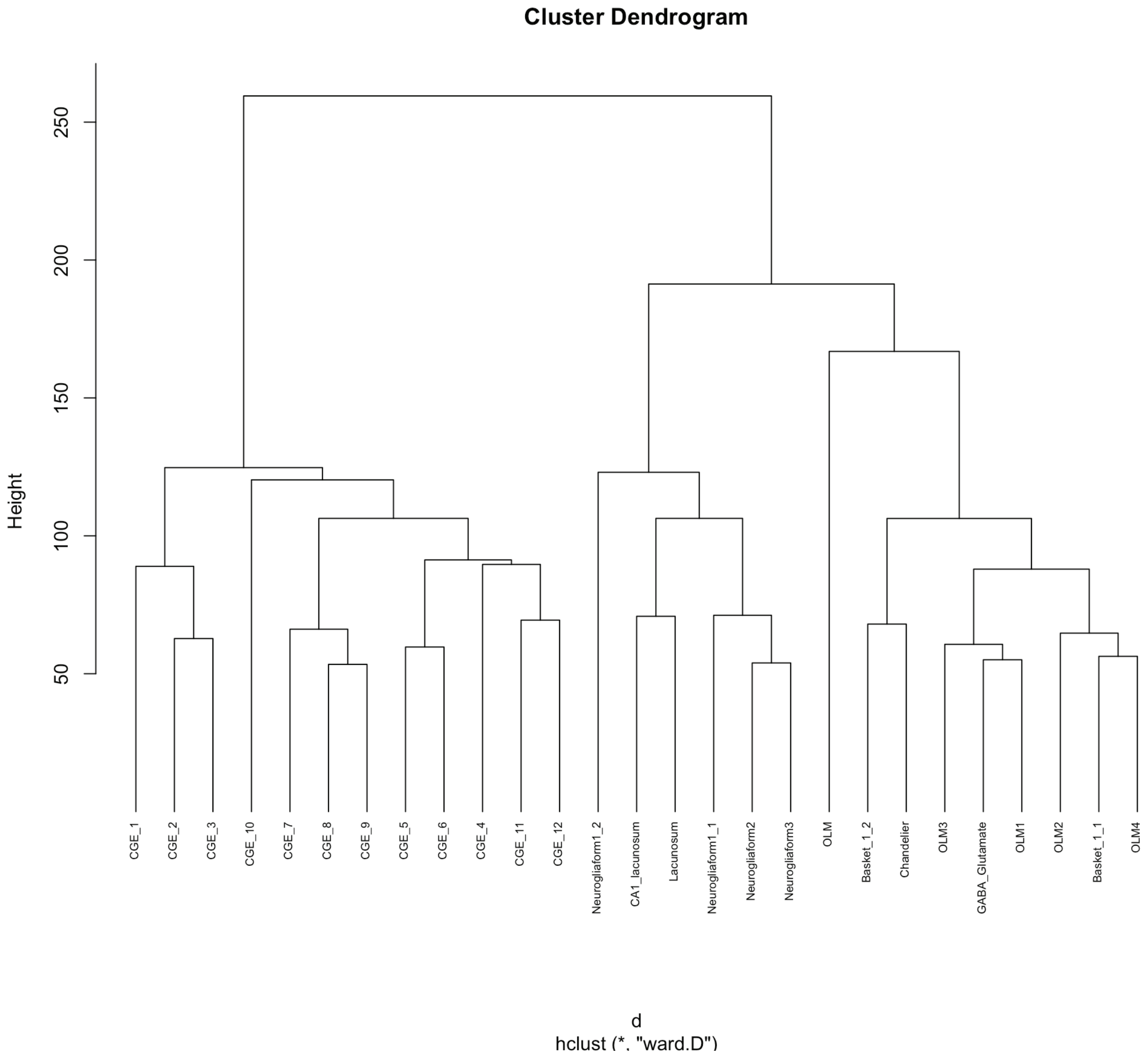
Cluster Dendrogram
Height
250
200
150
100
50
CGE_1
CGE_2
CGE_3
CGE_10
CGE_7
CGE_8
CGE_9
CGE_5
CGE_6
CGE_4
CGE_11
CGE_12
Neurogliaform1_2
CA1_lacunosum
Lacunosum
Neurogliaform1_1
Neurogliaform2
Neurogliaform3
OLM
Basket_1_2
Chandelier
OLM3
GABA_Glutamate
OLM1
OLM2
Basket_1_1
OLM4
d
hclust (*, "ward.D")

Supplementary Figure 9: Dendrogram of interneuron subtypes.

Dendrogram of 27 interneuron subtypes that appeared in the scRNA-seq hippocampus dataset, hierarchically clustered with Ward clustering.

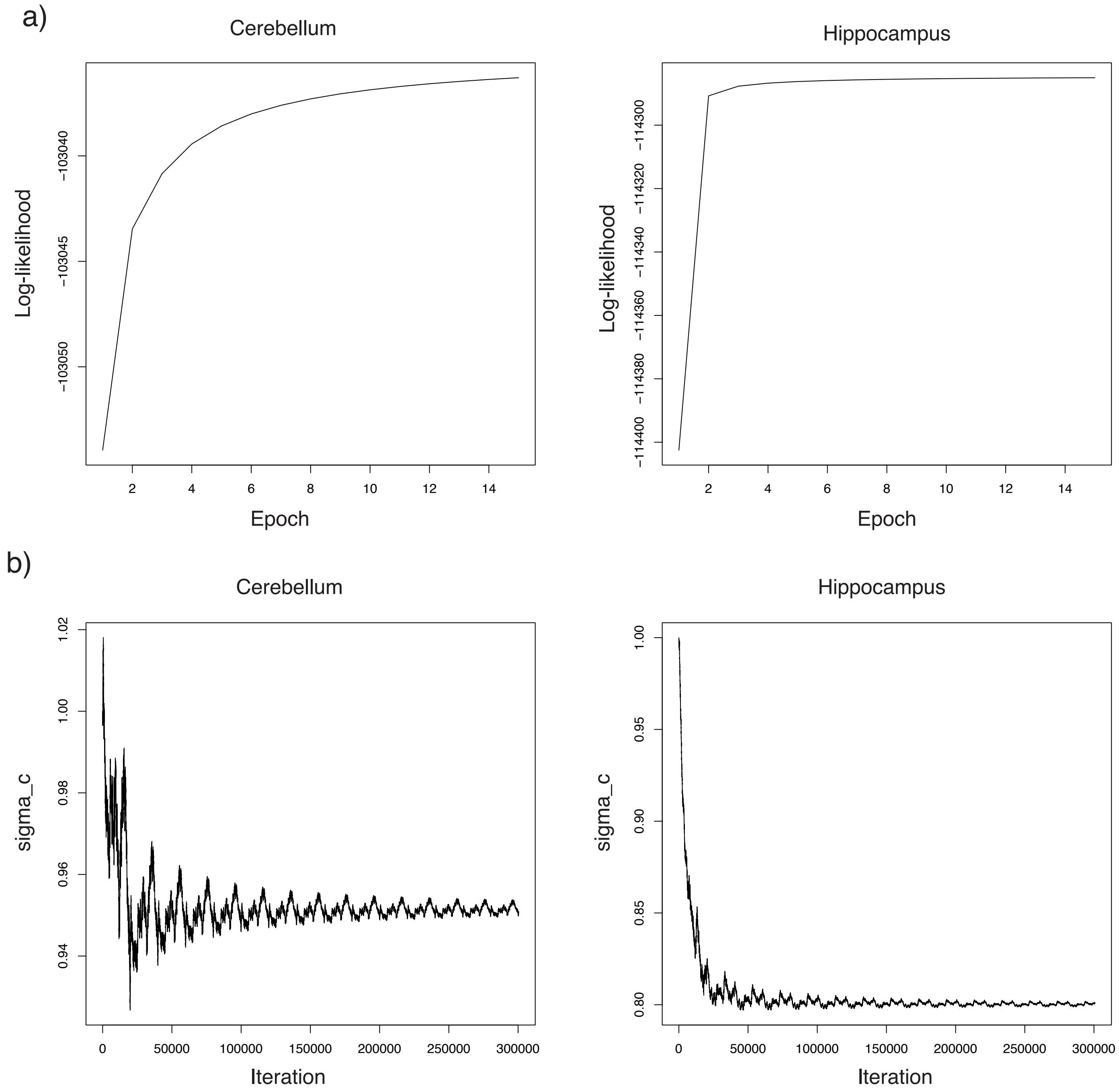
a)
Cerebellum
Log-likelihood
−103040
−103045
−103050
2
4
6
8
10
12
14
Epoch
Hippocampus
Log-likelihood
−114300
−114320
−114340
−114360
−114380
−114400
2
4
6
8
10
12
14
Epoch
b)
Cerebellum
sigma_c
1.02
1.00
0.98
0.96
0.94
0
50000
100000
150000
200000
250000
300000
Iteration
Hippocampus
sigma_c
1.00
0.95
0.90
0.85
0.80
0
50000
100000
150000
200000
250000
300000
Iteration

Supplementary Figure 10: Trace plots showing convergence of $\sigma_\varepsilon$ hyperparameter during optimization in RCTD.

a) Value of log-likelihood at each stochastic gradient descent (SGD) epoch optimizing $\sigma_\varepsilon$. Left: Slide-seq cerebellum. Right: Slide-seq hippocampus.

b) Value of $\sigma_\varepsilon$ at each stochastic gradient descent (SGD) iteration optimizing $\sigma_\varepsilon$. Left: Slide-seq cerebellum. Right: Slide-seq hippocampus.
